# Evolutionary dynamics of sex chromosomes and candidate sex-determining genes across the beetle phylogeny

**DOI:** 10.64898/2026.07.17.739015

**Authors:** Yuan Fu, Wenjing Sun, Bengt Hansson

## Abstract

The evolutionary dynamics of beetle sex chromosomes remain poorly understood. Using synteny analyses of 163 chromosome-level genome assemblies, we examine sex chromosome evolution across the beetle phylogeny. Beetles retain a highly conserved ancestral X chromosome region, even in superfamilies with extensive autosomal rearrangements and in species with enlarged neo-X chromosomes formed by fusions of autosomal segments. In contrast, Y chromosomes vary greatly in presence, size and gene content, driven by degeneration of ancestral regions and addition of neo-Y segments in some species. Neo-X and neo-Y chromosomes frequently occur and generally evolve independently, highlighting a flexible neo-sex chromosome dynamics. We identify putative sex-determining genes and propose a novel candidate, the highly conserved and sexually differentially expressed X-linked gene *ELAV-like*. Our results uncover conserved X elements, fragile Y chromosomes, and independent neo-X and neo-Y formation across 300 million years of beetle genome evolution.

## Introduction

Genetic sex determination via segregating sex chromosomes has evolved repeatedly across the tree of life from either hermaphroditic ancestors or systems with environmental sex determination (*1-4*). Although this process consistently produces separate males and females, the underlying genetic mechanisms and genomic architecture vary widely across taxa. For example, in therian mammals, a dominant Y-linked gene (*Sry*) triggers male development (*5*), whereas, in birds, sex determination depends on dosage of a Z-linked gene (*DMRT1*) (*6*). The complexity of the sex-specific region surrounding the sex-determining gene(s) also differ greatly: it is minimal in some fish and plants (*7-9*), but extensive in mammals and birds, where recombination cessation has led to heteromorphic sex chromosomes through progressive differentiation and degeneration (*10, 11*). Furthermore, sex chromosome dynamics range from extreme stability over millions of years (*9, 10*) to frequent transitions and turnovers, including shifts between male (XY) and female (ZW) heterogamety (*3, 12*). Understanding how this remarkable diversity in sex-determining mechanism and sex chromosome systems arises is a fundamental question, with broad implications for evolutionary processes such as sex-specific adaptation, sexual conflict and speciation (*13-18*).

Insects exhibit a wide range of sex-determining mechanisms and sex chromosome systems, including male and female heterogamety and haplodiploidy (*19, 20*). Sex determination in insects typically involves two key genes—*transformer* (*TRA*) and *doublesex* (*DSX*)—with *TRA* regulating sex-specific splicing of *DSX*, a transcription factor that directs male and female development (*21*). However, the initial signal triggering sex determination varies across taxa. The pathway is best characterised in *Drosophila*, where an X-linked polygenic region regulates sex determination through X-to-autosome balance (*22*). This balance activates *sex-lethal* (*SXL*) and *TRA*, which in turn mediate sex-specific splicing of *fruitless* (*FRU*) and *DSX* (*23, 24*). Other insects share similarities with the *Drosophila* pathway, but notable exceptions exist (*21, 25*), such as a male-determining factor in *Aedes* mosquitos (*26*) and a female-determining factor in *Bombyx* moths (*27*). Despite these insights, the evolutionary dynamics of sex determination and sex chromosome systems remain poorly understood for most insect lineages, including in beetles (Order Coleoptera), the most species-rich clade.

Coleoptera comprises over 350,000 described species, many exhibiting striking sexual dimorphism (*28*), and has diversified over the past 300 million years (*29-31*). Karyotype studies reveal haploid chromosome numbers ranging from 5 to 39 (commonly 9–12) and sex chromosome systems typically of the XO or XY type (*19, 32, 33*). A distinct sex chromosomal feature of several species if the parachute configuration, where a gene-poor Y chromosome (y_p_ or y_r_) does not recombine with X and primarily ensures correct segregation during male meiosis (*34, 35*). Complex systems are also common, including multiple X and/or Y chromosomes and neo-sex chromosomes formed by fusions of autosomal segments with the ancestral sex chromosomes (*32-36*). Despite the early discovery of a Y chromosome in the yellow mealworm beetle (*Tenebrio molitor*) in 1905 (*13, 17*), progress in understanding beetle sex determination has been limited. One of the few molecular studies, in the red flour beetle (*Tribolium castaneum*), suggests that maternal TRA protein initiates female development, while an unknown Y-linked gene may override this pathway to produce males (*37*). However, this mechanism cannot be universal, as many species have an XO system, suggesting that X-linked master regulators or lineage-specific sex-determining mechanisms may have evolved independently, as seen in Diptera (*25*). Recent genomic studies of individual beetle species (*38*) or small taxonomic samples (*39*) have characterised sex chromosomes and identified autosomes involved in neo-X chromosome formation (*39*). Nevertheless, a comprehensive comparative genomic analysis of sex chromosome dynamics and putative sex-determining genes across the beetle phylogeny remains lacking (*19, 40*).

Here, we analyse 163 publicly available chromosome-level beetle genome assemblies (*41*) spanning 39 families and 16 superfamilies to advance the understanding of sex chromosome evolution. Specifically, we (i) assess synteny conservation across autosomes and sex chromosomes, (ii) investigate the formation of enlarged neo-X and neo-Y chromosomes through autosomal fusions, identifying several previously uncharacterised neo-sex chromosomes, and (iii) examine the conservation and chromosomal location of key insect sex-determining genes to pinpoint candidate sex-determining genes in beetles.

## Results

### Highly conserved chromosome synteny across beetle superfamilies

To assess synteny across the beetle phylogeny, we selected one representative species from each of the 16 beetle superfamilies represented among our chromosome-level assemblies (Table S1), prioritising type genera when available (e.g., *Carabus problematicus*, Caraboidea). The reconstructed phylogeny (Fig. 1A), based on 515 single-copy gene sequences, was consistent with previously published species trees (*31*). As reference chromosomes, we used the eight autosomes and X chromosome of *Elmis aenea* (superfamily Byrrhoidea), which are highly conserved and correspond almost completely to the eight ancestral linkage groups (ALGs) recently reconstructed for the last common ancestor of beetles (*42*), except that two chromosomes in *E. aenea* are homologous to a single ALG. For simplicity, our *Elmis aenea* reference chromosomes are indexed by their colour in Fig. 1B (e.g., A_R_). Pairwise synteny block analysis between adjacent species in the phylogeny revealed extensive syntenic regions across most chromosomes (Fig. 1B). Nevertheless, as expected from the deep phylogenetic divergence, and the variation in genome size and chromosome numbers (Fig. 1A; Table S1), several intra-chromosomal rearrangements and chromosomal fissions and fusions were detected (Fig. 1B). For example, *Cantharis flavilabris* displayed one fission (involving A_Y_) and three fusions: two involving A_O_, A_G_ and parts of A_Y_, and one merging A_S_ and A_B_. In contrast, the X chromosome showed strong homology across the phylogeny, representing a highly conserved ancestral X region among beetles (Fig. 1B). This was confirmed by synteny analysis of RBH genes represented on the reference X chromosome, which further revealed that the ancestral X region varies roughly tenfold in size across species (Fig. 1C). Notably, five of the 16 species exhibited neo-X chromosomes formed by autosomal segments fused with the ancestral X (Fig. 1B).

**Fig. 1.**
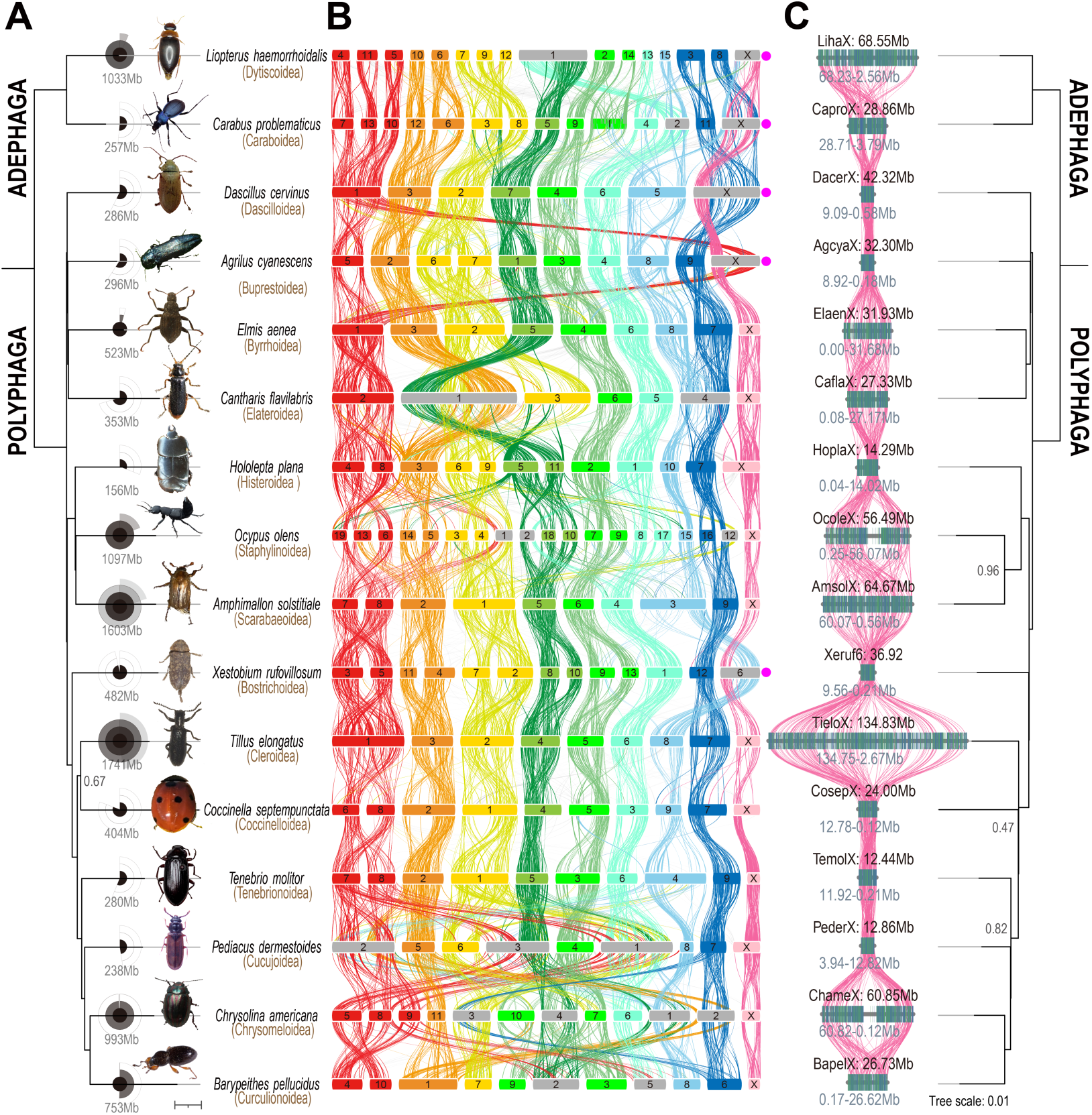
Chromosome evolution among beetles (16 superfamilies). (**A**) Phylogenetic species tree based on 515 single-copy genes (branch support = 1 unless indicated). Also shown are genome size (concentric rings: black = 0-0.5 Gb, dark gray = 0.5-1 Gb, gray = 1-1.5 Gb, light gray = 1.5-2 Gb), morphological representation of each species (images from https://www.gbif.org/), and species and superfamily name. (**B**) Pairwise synteny between adjacent species in the phylogeny. Links connect syntenic blocks and are coloured by homology to *Elmis aenea* reference chromosomes (autosomes: A_O_ orange, A_Y_ yellow, A_G_ green, A_L_ lime, A_C_ cyan, A_S_ skyblue, A_B_ blue; X chromosome in pink). Bars represent chromosomes scaled to genome size; colours indicate primary homology to *E. aenea* with gray bars indicating large fusion events. Filled pink dots mark neo-X chromosomes. (**C**) Synteny within the conserved ancestral X region. Links connect reciprocal best-hit (RBH) genes present in all species. Also, shown is a phylogenetic tree based on 171 X-linked RBH genes (branch support = 1 unless indicated).

### X chromosome stability also in superfamilies with scrambled autosomes

Synteny block analyses across 163 species revealed highly conserved syntenic patterns within most superfamilies (Fig. 2A–B; Figs. S1–S2; Table S2). However, four superfamilies— Caraboidea, Staphylinoidea, Curculionoidea and especially Chrysomeloidea—showed extensive autosomal rearrangement (Fig. 2B; Fig. S1). Within Chrysomeloidea, we identified four major clades with contrasting synteny: *Pogonocerus hispidulus*–*Rutpela maculata* exhibited well-conserved synteny; *Phaedon cochlieariae*–*Chrysolina americana* showed intermediate conservation; whereas *Crioceris asparagi*–*Cryptocephalus moraei* and *Altica lythri*–*Diorhabda carinata* displayed extensive rearrangements (Fig. 2B; Figs. S1 & S3). Even within some genera, multiple autosomal rearrangements occurred including both fissions and fusions (e.g., within *Diorhabda*, Chrysomeloidea; Fig. 2B; Figs. S1 & S3). Remarkably, the ancestral X region remained highly conserved even in superfamilies with otherwise highly scrambled autosomes (Fig. 2B; Figs. S1 & S4). In one species, *Holotrichia oblita* (Scarabaeoidea), the majority of the X chromosome was missing from the assembly, while the autosomes were represented and well-conserved (Fig. 2B; Fig. S1; Table S2).

**Fig. 2.**
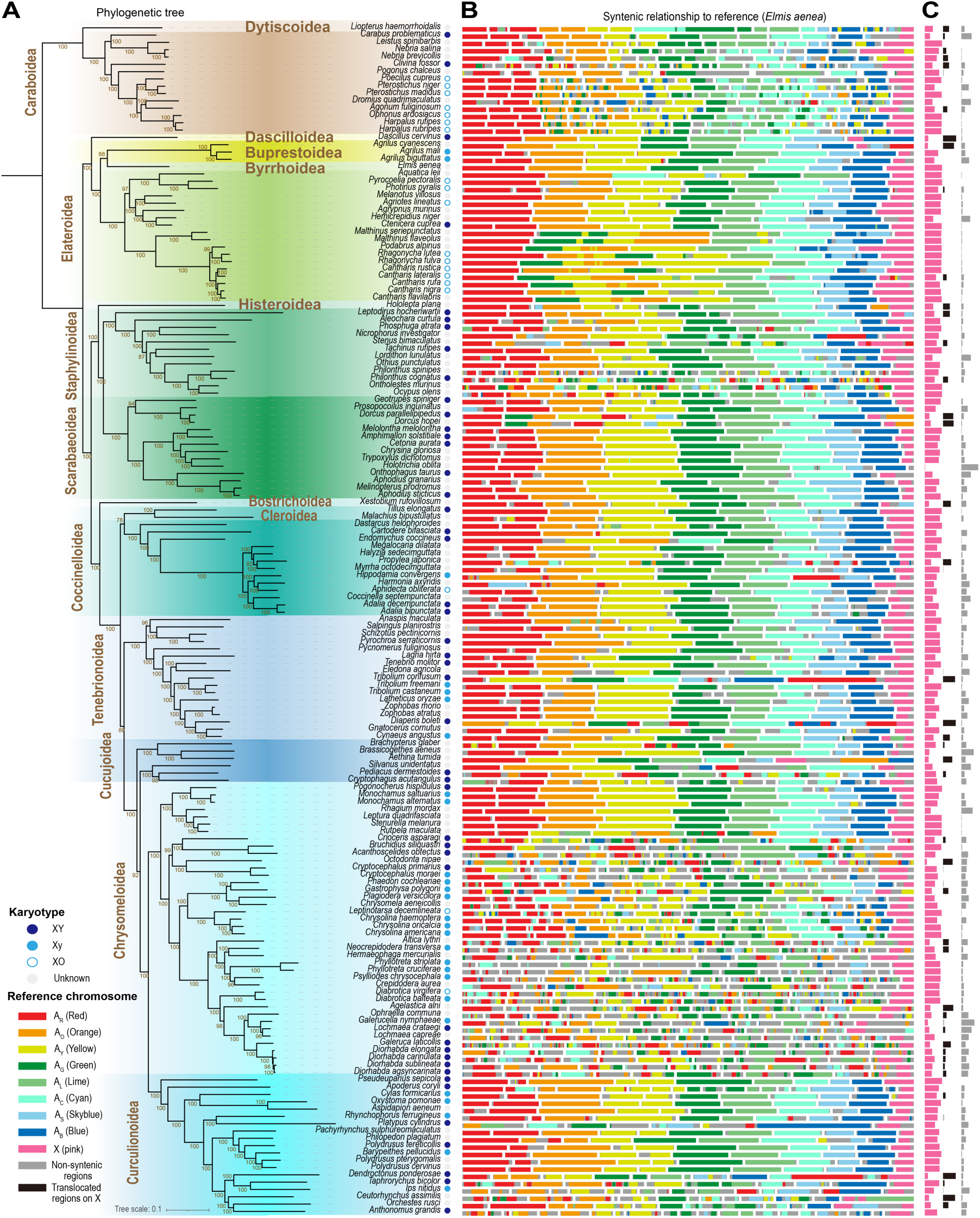
Phylogenetic relationship and chromosome evolution across 163 beetle species. (**A**) Phylogenetic species tree based on single-copy genes (branch support = 1 unless indicated). Also, shown are superfamily (gold), species (black) and sex chromosome system inferred from karyotypes (dots; (*32, 33*)). (**B**) Syntenic chromosome regions of each species relative to *Elmis aenea* reference chromosomes (coloured accordingly). Y chromosomes are excluded because most species lack assembled Y chromosomes. (**C**) Proportion of the X chromosome composed of conserved (pink), autosomal (black) and non-syntenic regions (grey).

### Neo-X chromosome formation involves every autosome

Neo-X chromosomes were detected in 51 of the 163 genomes, with segments from every autosome involved in at least two (A_C_) and up to eleven (A_R_) fusion events (Fig. 3A–C; Fig. S4; Table S2). Within species, one to five autosomes contributed to neo-X chromosome formation, the most extreme case being *Octodonta nipae* (Chrysomelidae), where parts of A_R_, A_O_, A_Y_, A_S_ and A_B_ were translocated to the ancestral X chromosome. Most rearrangements involved partial autosomal segments, but complete fusions of entire autosomes and X occurred in five species, including *Aethina tumida* (Cucujoidea) where A_O_ and A_B_ fused entirely to X. Among the 51 neo-X chromosomes, 21 showed no (post-fusion) rearrangement between the ancestral and added regions; of those with rearrangements, four belonged to the genus *Diorhabda* (Fig. 3A–C; Fig. S4). Synteny network cluster analysis of RBH genes confirmed the conserved ancestral X region and revealed several non-conserved clusters representing fused segments unique to individual species or shared among closely related taxa, particular within genera such as *Dorcus* (*D. parallelipipedus* and *D. hopei*) and *Diorhabda* (four species) (Fig. S5A).

**Fig. 3.**
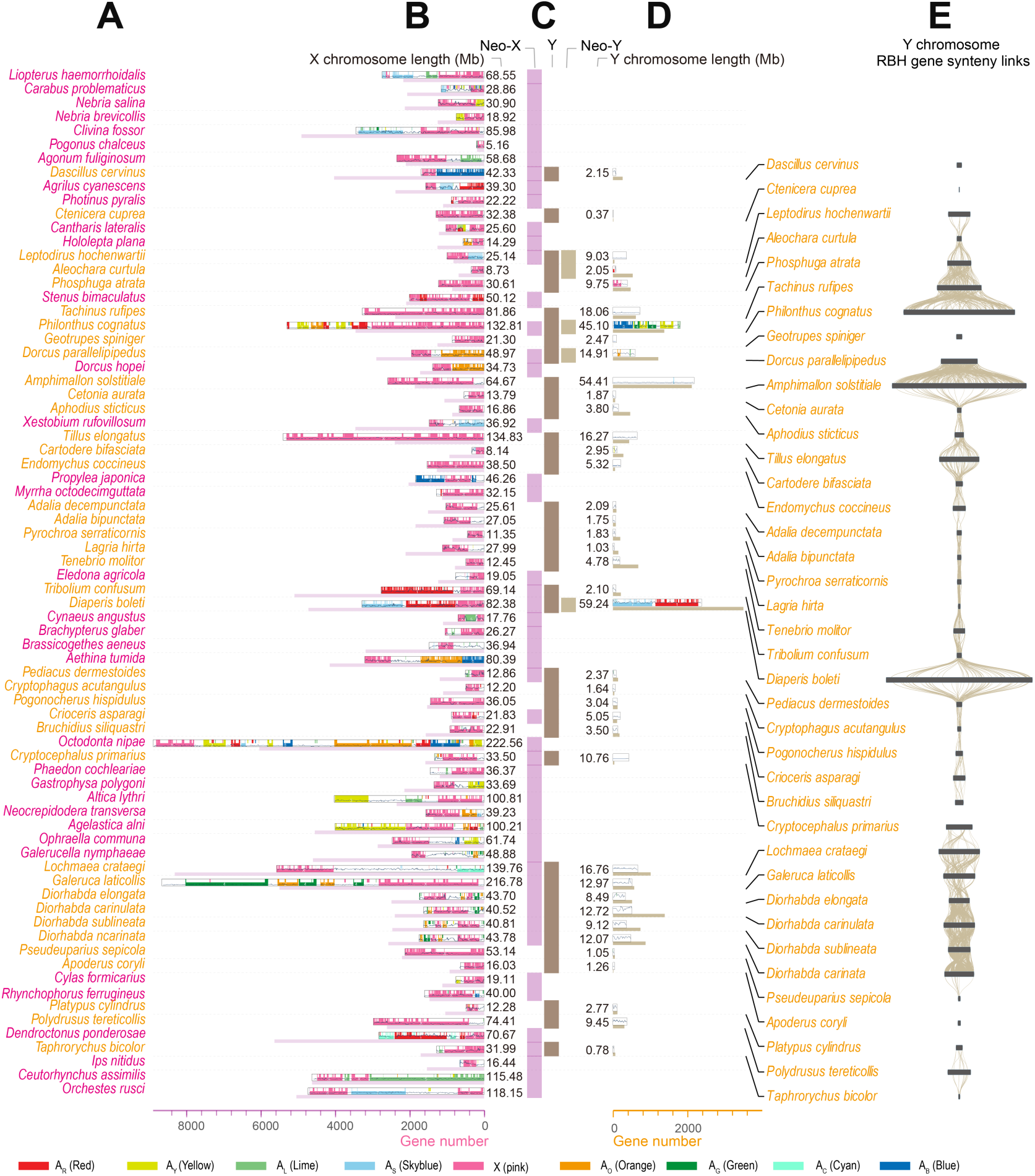
Syntenic regions, length and gene numbers of neo-X, Y and neo-Y chromosomes. (**A–D**) X and Y chromosome characteristics in species with neo-X chromosomes (51 species) and/or assembled Y/neo-Y chromosomes (39 species)—74 species in total—in phylogenic order. (**B,D**) X and Y chromosome length (box width), synteny to *Elmis aenea* reference chromosomes, gene density (line graph inside box), and gene count (horizontal bar; scale at bottom). (**C**) Species having neo-X (purple), Y (brown) and/or neo-Y chromosomes (light brown), respectively. (**E**) Synteny across Y chromosomes of 39 species. Links connect reciprocal best hit (RBH) genes between pairs of species.

Karyotype data were available for 21 of the 51 species with neo-X chromosomes, but only in five of those species had the neo-sex chromosome previously been described (*32, 33*). Thus, we identified 16 previously uncharacterized neo-X chromosomes among karyotyped species and 30 among species lacking karyotypes (Fig. 2B; Table S2).

Compared to autosomes, the X chromosome in most species contained large regions with poor gene content and no clear synteny to the reference genome (up to 74.8%; excluding *Holotrichia oblita* in which X is missing from the assembly), a pattern similar for neo-X regions (up to 75.1%) (Fig 2B–C; Figs. S1 & S4; Table S2). These gene-poor regions occurred almost always outside the conserved ancestral X region (Fig 2B–C; Figs. S1 & S4). Their origin of these gene-poor remains unclear, but they may represent centromeres, repeat-rich regions and/or translocated segments of degenerated Y chromosomes.

### Highly degenerated Y chromosomes and independently formed neo-Y chromosomes

The Y chromosome was present in 39 of the 163 genome assemblies (Fig. 3C; Table S2). The XO system is overrepresented in Caraboidea and Elatoroidea (*32, 33*), and among these clades only one of the 32 assemblies included a Y chromosome: *Ctenicera cuprea* (Elateroidea), which had a short (0.4 Mb) Y chromosome. Although no karyotype is available for this species, data from 19 other *Ctenicera* species show 12 with XO and seven with Xy_p_ karyotypes (*32, 33*). In the remaining 14 superfamilies, XY or Xy_p_ systems dominate (*32, 33*), yet only 38 of 131 assemblies contained a Y chromosome, suggesting that many of the assemblies are incomplete and lack Y sequences.

Synteny analysis against the *Elmis aenea* reference revealed neo-Y chromosomes in six species (Fig. 3C–D; Table S2). These formed through fusions of autosomal segments: A_R_ in *Aleochara curtula*; A_Y_, A_G_, A_L_ and A_B_ in *Philonthus cognatus*; A_O_ and A_C_ in *Dorcus parallelipipedus*; A_S_ in *Amphimallon solstitiale*; A_Y_ in *Cartodere bifasciata*, and the entire A_S_ and parts of A_R_ in *Diaperis boleti*. Karyotype data were available for three of these six species, in which neo-Y chromosomes were reported in two (*32, 33*). Thus, we identified four new neo-Y chromosomes; one among species with a karyotype and three among species without. Neo-Y chromosomes were generally larger (2.0 – 59.2 Mb) than other Y chromosomes (0.4 – 18.0 Mb), consistent with recent additions (Fig. 3D). Only *Diaperis boleti* showed homology between its neo-Y and neo-X chromosomes, both containing parts of A_R_ and the entire A_S_ (Fig. 3A–D). In contrast, other neo-Y chromosomes evolved fully or partly independently of their X chromosomes, which highlights a pronounced flexibility in neo-X and neo-Y formation across beetles. Note, however, that *Philonthus cognatus* may represent an assembly problem. This species appears to possess a complex neo-Y containing parts of A_Y_, A_G_, A_L_ and A_B_ and a neo-X formed by parts of A_R_, A_O_ and A_Y_ (Fig. 3A–D), a highly unusual configuration in beetles (see also (*42*)).

Because the *Elmis aenea* reference lacks a Y chromosome, potentially obscuring conserved regions, we performed a synteny analysis of RBH genes across the 39 species with a Y chromosome. Synteny links between the RBH genes were primarily observed among ancestral Y chromosomes of closely related species (e.g., *Lochmaea*, *Galeruca* and *Diorhabda*), whereas there was no evidence of conserved genes across the entire phylogeny (Fig. 3E). That neo-Y chromosomes lack syntenic relationships to other Y chromosomes reflects that different autosomal segments have fused to the Y chromosome in different lineages (Fig. 3D). A synteny network cluster analysis confirmed this patter, revealing no strongly conserved clusters but several scattered clusters representing rearrangements unique to individual species or small clades (Fig. S5B). The lack of conservation on the Y chromosome aligns with the prevalence of XO or Xy_p_ systems in beetles and mirrors patterns seen in other ancient sex-limited chromosomes, such as the mammalian Y and avian W, which are also highly degenerated (*10, 18, 43*).

Finally, the distal part of Y chromosome of *Phosphuga atrata* showed substantial synteny with the conserved ancestral X region (Fig. 3D; Table S2). This pattern could suggest recent additions of X-linked segments to the Y chromosome. However, because the corresponding genes are absent from its X chromosome (see below; Table S2), we interpret this as a misassembly.

### Conserved gene content on the X but not on the Y chromosome

The X chromosome length ranged between 5.16 and 222.56 Mb (median 32.96 Mb), and the gene count between 190 and 8,308 (median 1,504) across assemblies (Figs. 2B–3B; Fig. S4; Table S2). This variation largely reflects the presence of substantially enlarged neo-X chromosomes in some species (e.g., *Clivina fossor*, Caraboidea) and assembly errors or incompleteness in three species (see below; Table S2). To further explore variation in X-linked gene content, we analysed the number of species sharing specific orthogroups (OGs) represented by at least one X-linked gene copy (Fig. 4A; Table S3). Note that an OG can include multiple genes located on different elements (X, Y, autosomes or scaffolds). The X chromosome exhibited two distinct peaks in OG-sharing: one corresponding to neo-X regions unique to single species or small clades with shared autosomal translocations (left peak; Fig. 4A), and another representing the conserved ancestral X region shared by most species (right peak; Fig. 4A).

**Fig. 4.**
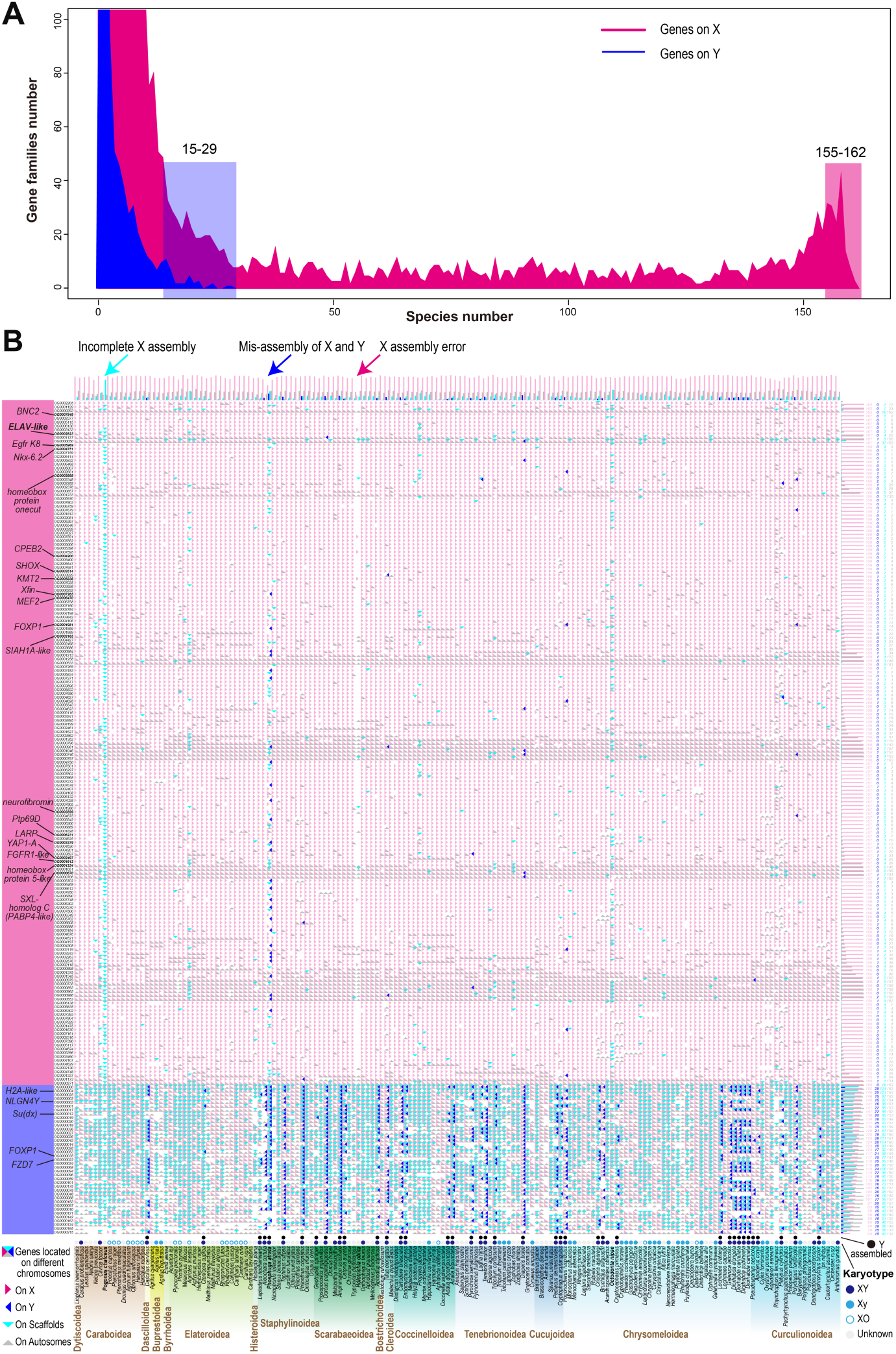
X-linked and Y-linked orthogroups (OGs) shared across species. (**A**) Distribution of OGs shared by different numbers of species for X-linked OGs (red; OGs including at least one X-linked gene) and Y-linked OGs (blue; OGs including at least one Y-linked gene). The X-linked distribution shows two peaks: one representing neo-X-linked OGs (left; OGs shared by species with the same autosomal addition) and another representing the conserved ancestral X region (right; OGs shared by nearly all species). The distribution of Y-linked OGs shows a single peak, representing neo-Y-linked OGs (left), and a tail representing OGs found in relatively many assemblies. Boxes highlight the most widely shared X-linked (red) and Y-linked (blue) OGs. (**B**) Chromosomal location of the most widely shared X-linked (red) and Y-linked (blue) OGs. Shown are OGs occurring on the X chromosome in ≥155 species or on the Y chromosome in ≥15 species. Each cell shows OGs with genes located on the X (pink), Y (blue), scaffolds (cyan), and/or autosomes (grey). OG index and names of putative sex-related genes are listed (left). Histograms (top) and numbers (right) summarize data per species (top) and OG (right). Species are phylogenetically ordered (Fig. 2A). Sex chromosome system inferred from karyotypes (coloured dots) and presence of an assembled Y chromosome (dots) are indicated above species names. Arrows highlight species with X assembly errors: *Pogonus chalceus* (cyan arrow; incomplete X assembly: conserved X-linked OGs on scaffolds), *Phosphuga atrata* (blue arrow; mis-assembly of X and Y: many conserved X-linked OGs only on Y chromosome), and *Holotrichia oblita* (red arrow; X chromosome partly missing: majority of conserved X-linked OGs missing).

We next examined the genomic location of all genes in OGs represented on the X chromosome in most species (≥155 species), totalling 179 OGs (Fig. 4B; Table S3). Most of these OGs were exclusively X-linked, although some were also present on autosomes. Only in a few species these OGs were represented on the Y chromosome and on unplaced scaffolds. OGs consistently found on both X and autosomes likely represent ancient gene duplication events. Three species exhibited exceptionally few X-linked OGs due to assembly errors or incompleteness: *Pogonus chalceus* (Caraboidea) has an incomplete X assembly with most OGs located on unplaced scaffolds (Fig. 4B); *Phosphuga atrata* (Staphylinoidea) shows a misassembly of X and Y (Fig. 4B; Fig. S6A; Table S2); and for *Holotrichia oblita* (Scarabaeoidea) the majority of the X chromosome is missing (Fig. 4B).

The length of Y and neo-Y ranged between 0.37 and 59.24 Mb (median length 3.80 Mb), and the gene count between 3–3,501 (median 211; Fig. 3C–D; Fig. S5C; Table S2). This variation primarily reflects the presence of neo-Y chromosomes in some species. Consistent with the lack of synteny of beetle Y chromosomes, we found little evidence of conserved gene content (Fig. 3E; Fig. S5B). Large, relatively gene-rich Y chromosomes occurred only in species with neo-Y chromosomes, whereas species without autosomal additions had short, gene-poor Y chromosomes (Fig. 3C–D; Fig. S5C). Analysis of OGs containing Y-linked genes reinforced this pattern (Fig. 4A–B). The overwhelming majority of Y-linked OGs were unique to single species or shared only by a few species (Fig. 4A). Very few Y-linked OGs were shared broadly, and none occurred in more than 29 of the 39 species with Y chromosomes (39 OGs present in ≥15 species are shown in Fig. 4B). Moreover, nearly all Y-linked OGs shared by multiple species included multi-copy genes also located on the X chromosome, autosomes or unplaced scaffolds (Fig. 4B), providing very weak support for conserved Y-linked genes.

### Candidate sex-determining genes

To identify candidate sex-determining genes in beetles, we first examined the functional annotation of the 179 X-linked and 39 Y-linked OGs overrepresented on their respective chromosome (Fig. 4B; Table S3). None of these are previously described sex-determining genes, but several share homology with such genes or have annotated sex-related functions in various organisms (X-linked: 17 OGs including *BNC2* (*44*), *ELAV-like* (*45*), *Egfr K8* (*46, 47*), *LARP* (*48*), *PABP4-like* (*48, 49*) and *YAP1-A* (*50*); Y-linked: 5 OGs including *H2a-like* (*51*) and *NLGN4Y* (*52*); Table S3). For example, *BNC2* (*basonuclin-2-like*) encodes a zinc-finger protein essential for proper mitotic arrest and meiotic progression in mouse male germ cells (*53*).

Next, we screened the genomes for homologs of key insect sex-determining genes (*20*)—*sex-lethal* (*SXL*), *transformer* (*TRA*), *fruitless* (*FRU*) and *doublesex* (*DSX*)—using moderately stringent matching criteria. Multiple homologs were detected in most genomes, located primarily on autosomes, with some on X chromosomes and none on Y chromosomes (Fig. 5A; Fig. S7). Gene tree analyses of these homologs showed no X-linked clusters for *TRA*, *FRU* and *DSX* (Fig. S7). In contrast, the gene tree of all *SXL* homologs revealed three clusters (*SXL-homolog A, B* and *C*), each highly conserved and predominantly X-linked (Fig. 5A). Reanchoring these homologs to eight well-annotated beetle genomes (*Aethina tumida*, *Anthonomus grandis*, *Coccinella septempunctata*, *Diabrotica virgifera*, *Diorhabda carinulata*, *Diorhabda sublineata*, *Harmonia axyridis* and *Tribolium castaneum*) showed that *SXL-homolog A* and *B* were co-localised on *ELAV-like* (*embryonic lethal abnormal vision-like protein 3*; Fig. 5B), while *SXL-homolog C* corresponds to *PABP4-like* (*polyadenylate-binding protein 4-like*). We reannotated *SXL*, *PABP4-like* and *ELAV-like* by using published protein sequences and constructed gene trees across all species (Fig. S8), as well as detailed comparisons among the eight reference assemblies (Fig. 5B). *SXL* was present in only 64 species and X-linked in just one, whereas *PABP4-like* was X-linked in 156 species and autosomal in 18. *ELAV-like* was X-linked in 161 of the 163 species, with only two species also carrying an autosomal copy (Fig. S8). Its absence in *Holotrichia oblita* reflects the partly missing X chromosome in the assembly, and in *Pogonus chalceus* it is located on an unplaced scaffold (Fig. 4B; Fig. S8D), consistent with X being incompletely assembled. Notably, *ELAV-like* exhibited the highest sequence conservation among these three genes (indicated by short branch lengths in the gene trees; Fig. 5B; Figs. S7– S8), suggesting strong purifying selection.

**Fig. 5.**
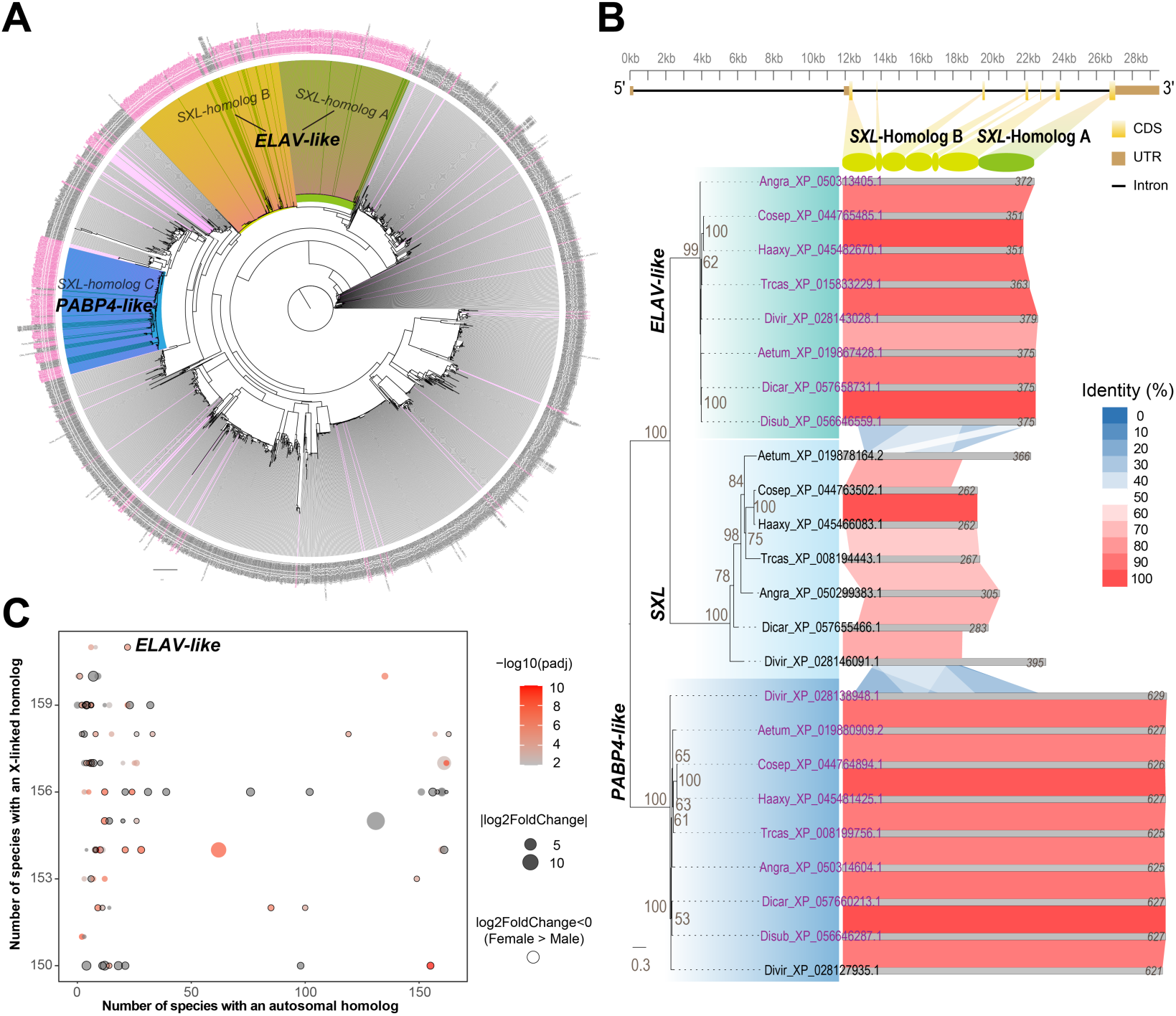
Gene-tree clustering of *SXL* homologs and sex-biased expression of conserved X-linked genes. (**A**) Gene tree of all *SXL* homologs across 163 species. The outer circle shows species names, with purple indicating X-linked homologs. (**B**) Gene tree of three *SXL* homologs—*ELAV-like*, *SXL* and *PABP4-like*— extracted from eight well-annotated genomes. Gene names in purple indicate X-linked copies. Top: gene structure of *ELAV-like* in *Anthonomus grandis* showing coding sequences (CDSs: yellow boxes), untranslated regions (UTRs: brown) and introns (black line). Next to the phylogeny: protein sequence length (narrow grey bars) and pair-wise synteny links, colour-coded by sequence identity (%). ELAV-like shows approximately 45% identity to the SXL protein. (**C**) Sex-biased expression among conserved X-linked genes in *Ips typographus*. Symbol sizes indicate degree of sex-biased expression (|log_2_FoldChange|), and colour indicates significance level (-log_10_ adjusted *p*-value). Each gene is plotted according to the number of species in which the gene has an X-linked homolog (y-axis) and an autosomal homolog (x-axis). *ELAV-like* is indicated at the top.

Finally, we downloaded and analysed available gene expression data from adult male and female *Ips typographus* (*54*). *ELAV-like* was among the few conserved X-linked genes with sex-biased expression, showing higher expression in females than in males (female-to-male expression ratio = 1.69; *P_adj_* = 1.15 × 10^-5^; Fig. 5C; Table S4).

Taken together, its consistent X-linkage, homology to *SXL*, high sequence conservation and female-biased expression make *ELAV-like* a compelling candidate for the primary sex-determining gene in beetles.

## Discussion

This comprehensive comparative study of 163 beetle genomes, spanning 16 superfamilies, represents a major advance in understanding the evolutionary dynamics of insect sex chromosomes (*35, 39, 55*). First, we show that a highly conserved ancestral X region has retained numerous X-linked genes over ∼300 Myrs of beetle evolution (*29-31*), consistent with previous and recent studies (*39, 42*). Second, we demonstrate that neo-X and neo-Y chromosomes frequently arise through fusions of one or more autosomal segments to the ancestral sex chromosomes, with all reference autosomes contributing multiple times to neo-sex chromosome formation. Third, we confirm that Y chromosomes are generally absent (XO systems) or small and gene-poor (XY and Xy_p_ systems) (*19, 32, 33*), except in species with neo-Y chromosomes, which contain variable numbers of recently translocated genes. Several of the neo-X and neo-Y chromosomes identified here have not been previously described. Finally, we show that *ELAV-like* shares homology with the well-known insect sex-determining gene *SXL*, is consistently X-linked, exhibits strong sequence conservation across the beetle phylogeny and is sexually differentially expressed, making it a strong candidate for primary sex determination in beetles.

Our analysis reveals a general pattern of extensive synteny across and within most superfamilies with large syntenic regions detectable on all chromosomes. Nevertheless, as expected from deep phylogenetic divergence and substantial variation in genome size and chromosome numbers, most homologous regions exhibit intra-chromosomal rearrangements, and chromosome fissions and fusions are common. Four superfamilies—Caraboidea, Staphylinoidea, Curculionoidea and especially Chrysomeloidea—deviate markedly from an organized, conserved pattern, displaying highly rearranged autosomal genomes. Remarkably, even in these superfamilies with highly rearranged genomes, the ancestral X region remains strongly conserved. This finding in beetles parallels observations in true bugs (Hemiptera), where highly conserved X chromosomes coexist with extensively rearranged autosomes (*56, 57*). Interestingly, this pattern goes outside insects, as mammals were recently shown to have a large, highly conserved X-linked region (*58*).

We confirm several previously reported neo-sex chromosomes (e.g., in *Tribolium confusum* (*39*) and *Dendroctonus ponderosae* (*38*)) and uncover numerous previously undescribed cases of neo-XY, Xneo-Y and neo-Xneo-Y systems (e.g., in *Dascillus cervinus* (*59*), *Dorcus hopei* (*60*)). Notably, most neo-X and neo-Y chromosomes appear to have evolved fully or partly independently, rarely sharing fused autosomal segments within species. Such independence may help explain the emergence of highly complex multiple-X and -Y karyotypes observed in some beetles. For example, the X_1_X_2_X_3_X_4_Y_1_Y_2_ system of *Blaps tenuicollis* (Tenebrionoidea) (*32, 33*) may have arisen through multiple fissions or separate autosomal fusions to X and Y. Unfortunately, this hypothesis cannot be directly tested because no chromosome-level genome assembly exists for this species, and no comparably complex karyotypes were present among species available for our study (Tables S1 & S2) (*32, 33*). In the related *Blaps rhynchoptera* (not included in our comparative study), newly released assembly data suggest that its X_1_X_2_Y system appears to result from a fission of the ancestral X (*61*) (NCBI accession number ASM4225770v1; Fig. S9).

Synteny analyses depend on genome assembly quality. Most assemblies analysed here were generated using long-read sequencing combined with HiC scaffolding (*62*), suggesting overall high quality. However, some genomes appear incomplete or misassembled. First, the majority of the X chromosome in *Holotrichia oblita* (Scarabaeoidea) is missing—an unlikely scenario without severe physiological and reproductive consequences. Most parts of the short, gene-poor chromosome labelled X may instead represent a degenerated Y or y_p_ chromosome. Karyotype data indicate that three other *Holotrichia* species have Xy_p_ systems, and most Scarabaeoidea species are XY or Xy_p_ (*19, 32, 33*). Second, another misassembly involves *Phosphuga atrata* (Staphylinoidea), which uniquely shows a Y containing many genes typically located on X. While an X-to-Y translocation is theoretically possible, its Xy_p_ karyotype (*32, 33*) suggests a degenerated Y, and moreover loss of numerous X-linked genes would render them absent in females (XX), causing severe functional consequences. Finally, the X chromosome of *Pogonus chalceus* (Caraboidea) is incomplete, as most ancestral X-linked genes—including *ELAV-like—*are located on unplaced scaffolds, indicating a need for further scaffolding. Non-recombining sex chromosomes are often highly degenerated and repeat-rich, making them difficult to assemble and potentially incomplete even when the remaining chromosomes are accurately reconstructed.

A primary sex-determining gene in beetles has not yet been identified. In our screening for homologs of key insect sex-determining genes (*SXL, TRA*, *FRU* and *DSX*) (*21*), most were located on autosomes. However, among the *SXL* homologs there are two notable exceptions: *PABP4-like*, which is X-linked in nearly all species (n = 156), and *ELAV-like*, which is X-linked in all 161 species with complete assemblies (absent only in *Holotrichia oblita*, where the X is partly missing, and in *Pogonus chalceus*, where the X assembly is incomplete and *ELAV-like* resides on an unplaced scaffold). *ELAV-like* meets key criteria for a candidate sex-determining gene: homology to *SXL*, strong sequence conservation, consistent X-linkage, and sexually differential expression. The presence of an X-linked factor suggests that sex determination in beetles may depend on the X-to-autosome ratio (X:A) as in *Drosophila* (*22*). Surprisingly few detailed studies of beetle sex determination exist, with the most comprehensive in the red flour beetle (*Tribolium castaneum*). In this species, maternal TRA protein triggers female development, while an unknown Y-linked gene was hypothesised to override this mechanism to produce males (*37*). However, among the 39 assembled Y chromosome, we found no consistently conserved candidate genes. Y-linked genes occurring in multiple species (≥15 species), including some with potential sex-related functions (Table S3), belong to gene-rich OGs also present on autosomes, the X chromosome and/or on scaffolds. Thus, a male-determining mechanism based on X:A ratio rather than Y-linkage appears more likely across beetles. This interpretation does not contradict the findings in *Tribolium castaneum* (*37*), and accommodates XO systems, which are common occurring in beetles (*19, 32, 33*).

The emergence of an unexpected candidate gene may explain why the primary sex-determining gene in beetles has remained elusive. In *Drosophila*, *ELAV* does not function in sex determination (*63, 64*) (only one copy is X-linked), but it contains three RNA recognition motif (RRM) domains (RRM1/2/3) that show strong similarity to the SXL protein (RRM1/2) (*65*). If *ELAV-like* has adopted a sex-determining role in beetles, this would represent a novel function— an evolutionary shift that, while unexpected, is not unprecedented in insects. For example, in the yellow fever mosquito (*Aedes aegypti*), *Nix* has taken the primary sex-determining role (*26*), and a similar evolutionary event could have occurred ancestrally in beetles. In *Drosophila*, the early sex determination is controlled by the X-signal element (XSE), an X-linked polygenic region including *Scute*, *SsiA*, *Runt* and *Upd* (*20*). We do not exclude the possibility that multiple X-linked genes act together to regulate sex development in beetles. Indeed, the long-term conservation of a gene-rich ancestral X region supports this scenario. In addition to *ELAV-like*, we identified other well-conserved X-linked genes with putative sex-related functions, notably *PABP4-like* (*49*) (X-linked in 156 species) and *BNC2* (*66-68*) (X-linked in 161 species). Therefore, our suggestion of specific X-linked sex-determining gene(s) should be regarded as a hypothesis requiring experimental validation. Moreover, our analysis does not encompass all sex-determining systems in beetles. Several weevils and bark beetles (Curculionoidea) exhibit haplodiploid sex determination, where conventional sex chromosomes are absent (*19, 69*). In haplodiploid honeybees (Hymenoptera), female development depends on heterozygosity at the gene *CSD*, while hemi- or homozygosity leads to male development (*70*). A similar mechanism could involve a gene within the conserved ancestral X region in haplodiploid beetles. Exciting opportunities remain to unravel sex-determining processes in beetles, and our study provides X-linked candidate genes for future research, particularly *ELAV-like*.

## Materials and Methods

### Chromosome-level beetle genomes

We obtained chromosome-level genome assemblies for 163 beetle species, representing 16 Coleopteran superfamilies, from NCBI (accessed April 2024; Table S1). Most assemblies were generated using a combination of long-read sequencing and Hi-C scaffolding (see, e.g. (*71*)), but many lacked comprehensive gene annotation. In 45 assemblies, the X chromosome was not previously identified. Many assemblies lacked a Y chromosome, likely due to sequencing a female or an XO system (Table S2).

### Synteny analysis

The scale of the dataset—163 chromosome-level genomes spanning deep evolutionary divergence and with inconsistent annotation—presented a significant computational challenge for synteny analysis. To address this, we developed a computationally efficient genome- and gene-guided synteny analysis pipeline optimised for large-scale, superfamily-level genome comparisons (*AgSyn* v0.6.1; GitHub: https://github.com/abysw/agsyn). The pipeline integrates widely used tools, including *ORFfinder* (*72*), *mafft*–*trimAl*–*iqtree*–*ASTRAL* (*73-76*), *JCVI* (*77*) and *genewise2* (*78*). Its output files also enable additional downstream analyses, including synteny analyses with *WGDI* (*79*) and synteny network cluster analysis using *Infomap* (v2.7.1) (80). Briefly, we identified open reading frames (ORFs) in each genome, mapped homologous genes using publicly available protein sequences (see below), and used these high-confidence gene locations as anchors to define syntenic blocks and chromosome homology across species. The analyses are outlined below and in Fig. S10. Analyses and parameters were finalized after two rounds of testing and evaluation. The most computationally intensive part of the pipeline— the annotation, corresponding to steps 1-5 in Fig. S10—processes ∼0.5 Gb per CPU-hour (Fig. S11).

Initial runs showed that the chromosomes of *Elmis aenea* (superfamily Byrrhoidea), comprising eight autosomes and the X chromosome, were highly conserved and therefore suitable as reference chromosomes across the phylogeny (Figs. 1–2). This was further supported by their close correspondence to the eight ancestral linkage groups (ALGs) recently reconstructed for the last common ancestor of beetles, except that two autosomes in *Elmis aenea* are homologous to a single ALG (*42*). For clarity, we indexed the *Elmis aenea* reference autosomes by their colour in Figs. 1–2: A_R_, red; A_O_, orange; A_Y_, yellow; A_G_, green; A_L_, light green; A_C_, cyan; A_S_, skyblue, A_B_, blue. The X chromosome is referred to as X and shown in pink.

The initial runs also showed that the X chromosome had been incorrectly identified as an autosome in two assemblies, *Diorhabda carinulata* and *Aleochara curtula*; these annotations were corrected in this study (Tables S1–S2). Moreover, *Cryptophagus acutangulus* was previously suggested to possess two X chromosomes, X1 and X2, based on synteny analysis However, our analyses support only X2 as an X chromosome, whereas X1, which is homologous to A_R_, should be reclassified as an autosome; this annotation was therefore corrected in this study (Table S1–S2).

### Input genome assemblies and reference protein sequences (pipeline steps 1 and 4; Fig. S10)

Chromosome identifiers of each genome were standardised to conventional nomenclature (1, 2, 3, X, Y, etc.) (step 1). To compile a set of homologous proteins for gene evidence (step 4), we retrieved protein sequences from five well-annotated beetle genomes (*Anthonomus grandis*, GCF_022605725.1; *Diorhabda carinulata*, GCF_026250575.1; *Ophraella communa*, GCA_035357415.1; *Coccinella septempunctata*, GCF_907165205.1; *Tribolium castaneum*, GCF_000002335.3) and *Drosophila melanogaster*, dmel_r6.55. Redundant sequences were removed using *CD-HIT* (v 4.8.1) (*82*), retaining only unique proteins.

### Annotate high-confidence conserved ORFs as “genes” (pipeline steps 2, 3 and 5)

Simple repeats were marked using *TANTAN* (v49) (*83*), and the genome assemblies were fragmented into ∼2 Mb contigs by introducing breaks at these repeat regions (step 2). Open reading frames (ORFs) were predicted using *ORF-FINDER* (v0.4.3) (*84*) (step 3) and aligned against the unique protein set compiled in step 4 using *DIAMOND* (v2.1.8.162) (*85*). ORFs lacking significant similarity to any homologous protein were removed. Multiple ORFs mapping to the same gene location were merged into a single element, which were defined as a “gene”, with constituent ORFs treated as individual CDS regions. The resulting gene annotations were re-anchored to the chromosome-level assemblies (Step 5).

Final annotations were saved in GFF3 format (*Genus_species.gff*), specifying genomic locations of these conserved elements. Each element includes a single “gene” and a single “mRNA” feature, which may contain multiple “CDS” features representing different ORFs at that locus. Corresponding nucleotide (*Genes_species.cds*) and amino acid (*Genes_species.pep*) sequences were extracted from the GFF3 and genome FASTA files using *gffread* (v0.12.7) (*86*). Because individual CDS sequences correspond to single ORFs, the resulting protein sequences contained internal stop codons (represented as dots), which were removed in subsequent step (below).

### Identifying orthogroups and constructing species tree (pipeline steps 6–7)

After removing all internal stop codons, the protein sequences annotated in step 5 were used to identify orthogroups (OGs) with *OrthoFinder* (v2.5.5) (*87*) (step 6). From these, 515 single-copy genes were extracted to construct a species tree comprising all 163 beetle species, and an additional set of 1,889 single-copy genes was used to build a tree representing one species from each of the 16 superfamilies. Multiple-species sequence alignments for each single-copy gene were generated using *MAFFT* (v7.520) (*73*), and poorly conserved regions were trimmed with *TrimAl* (v1.4.rev15) (*74*). Gene trees were inferred using *IQ-TREE* (v2.2.6) (*75*), and a coalescent-based species tree was constructed from all gene trees using *ASTRAL* (v5.7.8) (*76*) (step 7; Figs. 1A–2A).

### Synteny analysis of species representatives of the 16 superfamilies (pipeline step 8)

One representative species was selected from each of the 16 superfamilies, prioritising species from the type genus where available (e.g., *Carabus problematicus* for Caraboidea). Pairwise syntenic relationships among species in the phylogeny were identified using *JCVI* (v 1.3.9) (*88*) with default parameters to define synteny blocks. Chromosomes were coloured according to the reference chromosomes of *Elmis aenea* (see above; Fig. 1B–2B).

Next, to assess the evolutionary conservation of the X chromosome across the 16 superfamilies, we compared the X chromosome of *Elmis aenea* to those of the other 15 species using *JCVI* with parameters *--full -n 1*. This analysis identified reciprocal best-hit (RBH) gene pairs; 171 RBH genes located on the X chromosomes in all 16 species were retained. A phylogenetic tree based on these genes was reconstructed following the approach described in step 7 (Fig. 1C).

### Synteny analysis across the 163 genomes (pipeline step 8)

Syntenic relationships between *Elmis aenea* and each of the other 162 species were identified using *JCVI* (v 1.3.9) (*88*) with default parameters to define synteny blocks. To visualize these relationships across the entire dataset, we developed a new file format (*.agsyn*) that integrates four key data components: standardized genome file (step 1), gene annotations (step 5), species order list (derived from the phylogenetic tree in step 7), and synteny blocks (step 8; function *agsyn.syn*). The visualization displays all genomes at uniform total length, with chromosomal segments coloured according to their homology to the reference chromosomes of *Elmis aenea*. Each chromosome plot is divided into two regions: the upper portion shows synteny blocks between the query and the reference, while the lower portion shows inferred homologous blocks, defined by collinear blocks of single reference chromosomes spanning ≥2 Mb. Additionally, original chromosome numbers are shown in black to the right of each chromosome, and the position of homologous genes are displayed above the chromosomes (generated in step 10, below). The complete visualisation is shown in Figs. S1A–E & S4, while parts of data are shown in Fig. 2B.

### Analysis of Y chromosome synteny links (pipeline step 8)

Syntenic relationships among Y chromosomes of 39 species were analysed by identifying RBH genes using *JCVI* (*88*) with the parameter *--full -n 1*. To enhance visualization clarity, an alpha value of 0.05 was applied to synteny link diagrams (Fig. 3E).

### Analysis of sex chromosome-specific orthogroups (pipeline step 9)

To identify conserved X- and Y-linked genes, we quantified for each orthogroup (OG) obtained from step 6 the number of genes located on the X chromosome, Y chromosome, unplaced scaffolds and autosomes (function *agsyn.sex*; step 9). The results were visualized per OG and species as a square divided into four coloured triangular compartments, indicating gene presence on X (pink), Y (blue), unplaced scaffolds (cyan) and autosomes (grey). In the main text, we focused on OGs with gene presence on the X chromosome in ≥155 species (≥95% of 163 species) or on the Y chromosomes in ≥15 species (noting that only 39 assemblies contained an assembled Y chromosome) (Fig. 4A–B; all OGs are listed in Table S3).

OG function annotations were derived from their genomic positions in two well-annotated beetle reference genomes (*Aethina tumida* and *Diorhabda carinulata*; function *agsyn.agsx.func*; Table S3).

### Identifying putative sex-determining genes (pipeline step 10)

To identify candidate sex-determining genes across beetle genomes, we searched for homologs of four key insect sex-determining genes: *sex-lethal* (*SXL*), *transformer* (*TRA*), *fruitless* (*FRU*) and *doublesex* (*DSX*). For this purpose, we used the function *agsyn.gwise2*, which implements a homolog annotation pipeline based on *GeneWise2* (wise2-4-1) (*78*), building upon a previously established framework (*89*). This analysis (step 10) involves (i) preparing the genomes and unique protein sequences of each gene, (ii) reducing redundancy using *CD-HIT* (v 4.8.1) (*82*), (iii) identifying protein-DNA matches using *TBLASTN* (package *BLAST* 2.14.1) (*90*), (iv) refining matches with *SOLAR* (v0.0.19) (*91*), and retaining only those with high alignment scores and meaningful overlap, (v) modelling gene structures for the curated regions using *GeneWise2*, (vi) generating a gene annotation file in GFF3 format, and (vii) extracting the CDS and protein sequences using *gffread* (*86*) (as in step 5).

### Building gene trees of key insect sex-determining genes (pipeline step 11)

We constructed gene trees for all homologs of each of the four key insect sex-determining genes (*SXL*, *TRA*, *FRU* and *DSX*) to evaluate sequence conservation (short branch lengths indicate high conservation) and to distinguish the genomic location of homologs (X-linked or autosomal; no homologs were found on Y chromosomes) (step 11). For each gene, homologs were aligned using *MAFFT* (*73*) (*muscle* (*92*) can also be used) and poorly conserved regions were removed from the alignments using *TrimAl* (*74*). Gene trees were then constructed with *IQ-TREE* (*75*) (Fig. 5A; Figs. S7).

### Analysis of three X-linked clades of *SXL* homologs

The gene tree of *SXL* homologs (from step 11) revealed three clades (designated A, B and C) that predominantly included sequences located on the X chromosome (Fig. 5A). To investigate these clades further, we traced the *SXL* homologs to eight well-annotated reference genomes (*Anthonomus grandis*, *Coccinella septempunctata*, *Diabrotica virgifera*, *Diorhabda carinulata*, *Diorhabda sublineata*, *Harmonia axyridis*, *Tribolium castaneum*). We found that clades A and B correspond to the same gene region, the *ELAV-like* gene (Fig. 5B), whereas clade C maps to a different region, the *PABP4-like* gene.

Using the function *agsyn.gwise2* (step 10), we reannotated *ELAV-like*, *PABP4-like* and *SXL* across all 163 genomes, using the proteins from the eight reference genomes. Homologs of these genes were filtered with *BLASTP* (e-value ≤1e-120), and gene trees were reconstructed following the same approach as above (step 11; Fig. S8).

### Confirming the X chromosome in each species (pipeline step 12)

By integrating the results from the synteny analysis (step 8) and the analysis of X-linked OGs (step 9), we confirmed presence of the X chromosome in all but two species. In *Holotrichia oblita*, the putative X chromosome showed <3% synteny to the reference and lacked most X-linked OGs (the low level of synteny suggests that the assembled chromosome may correspond to a Y chromosome). In *Pogonus chalceus*, most X-linked OGs (including *ELAV-like*) were located on an unplaced scaffold rather than on a designated chromosome (suggesting incomplete assembly).

### Synteny network cluster analysis of genes on X and Y chromosomes (step 13 in Fig. S10)

To illustrate the relationships among short sequences and/or multiple gene families, we employed a network-based approach that visualises both conserved and non-conserved genes, as well as homologous and non-homologous connections. Reciprocal best-hit (RBH) genes among all species were identified using *JCVI* (parameter: *--full -n 1*) (*88*), and synteny network cluster relationships among genes and species were calculated and visualised using *Infomap* (v2.7.1) (*80*). Separate networks were generated for genes located on X and Y chromosomes (Fig. S5A– B).

### Gene expression analysis in *Ips typographus*

RNA-seq data of adult *Ips typographus* were downloaded from NCBI (accession number PRJNA934749). The dataset included four biological replicates of each sex from the control group of an acetone-exposure experiment (*54*). Sample quality was assessed by principal component analysis (PCA). One biological replicate from each sex clustered separately from the remaining replicates and was therefore excluded from downstream analyses, leaving three females (FA1, FA2, and FA4) and three males (MA1, MA3, and MA5) for expression analysis.

Raw paired-end reads were quality filtered using *Trimmomatic* (v0.39) (*93*) with the parameters SLIDINGWINDOW:4:20, LEADING:20, TRAILING:20, and MINLEN:36 using the TruSeq2 paired-end adapter set. Filtered reads were aligned to the *Ips nitidus* reference genome (NCBI GCA_018691245.2) using *HISAT2* (v2.2.2) (*94*). Genome indices were generated using splice-site and exon information extracted from the reference gene annotation to improve splice-aware alignment. Reads with mapping quality ≥10 were retained, and alignments were sorted and indexed using *SAMtools* (v1.22.1) (*95*).

Gene expression was quantified using *StringTie* (v3.0.3) (*96*) with the reference annotation in expression-estimation mode (-e) and gene-level read counts were generated using the accompanying *prepDE.py* script. Differential gene expression between females and males was assessed using *DESeq2* (v1.42.1) (*97*). Genome-wide expression estimates for all annotated genes are provided in Table S4. The main analysis focused on genes located on the conserved X chromosome and identified as X-linked in ≥150 species.

## Acknowledgments

Bioinformatics analyses were performed on computational infrastructure provided by the National Academic Infrastructure for Supercomputing in Sweden (NAISS) at the KTH Royal Institute of Technology. We thank the Darwin Tree of Life, Wellcome Sanger Institute, for access to chromosome-level genomes, and Dmitry Filatov for hosting YF and WS at the University of Oxford, where part of this work was conducted.

## Funding

Swedish Research Council grant 2022-04996 (BH)

China Scholarship Council grant CSC202304910565 (YF)

China Scholarship Council grant CSC202404910491 (WS)

## Author contributions

Conceptualization: BH, YF

Methodology: YF, WS, BH

Investigation: YF, WS, BH

Visualization: YF, WS

Funding acquisition: BH

Project administration: BH, YF

Supervision: BH

Writing – original draft: BH, YF Writing – review & editing: BH, YF

## Competing interests

Authors declare that they have no competing interests.

## Data, code, and materials availability

All sequence data used in this study are publicly available on NCBI (accession numbers are provided in table S1). All annotated code, data and results are available at GitHub: https://github.com/abysw/agsyn.

## Notes

### Competing Interest Statement

The authors have declared no competing interest.

