## Supplementary Figures S1-S11 for "Evolutionary dynamics of sex chromosomes and candidate sex-determining genes across the beetle phylogeny"

**The PDF file includes:**

Figs. S1 to S11

**Fig. S1.**

**Chromosome evolution across 163 beetle species.** Syntenic chromosome regions of each species relative to *Elmis aenea* reference chromosomes. Each main bar represents a chromosome with the upper part showing syntenic blocks and the lower part the inferred reference chromosome regions (coloured by homology to the reference chromosomes). Numbers refer to source chromosome name. Y chromosomes are excluded because most species lack assembled Y chromosomes. Positions of homologs of five key sex-determining genes (SDG homologs) are indicated above the bars by triangles.

**(S1A) Superfamilies Dytiscoidea – Histeroidea:**

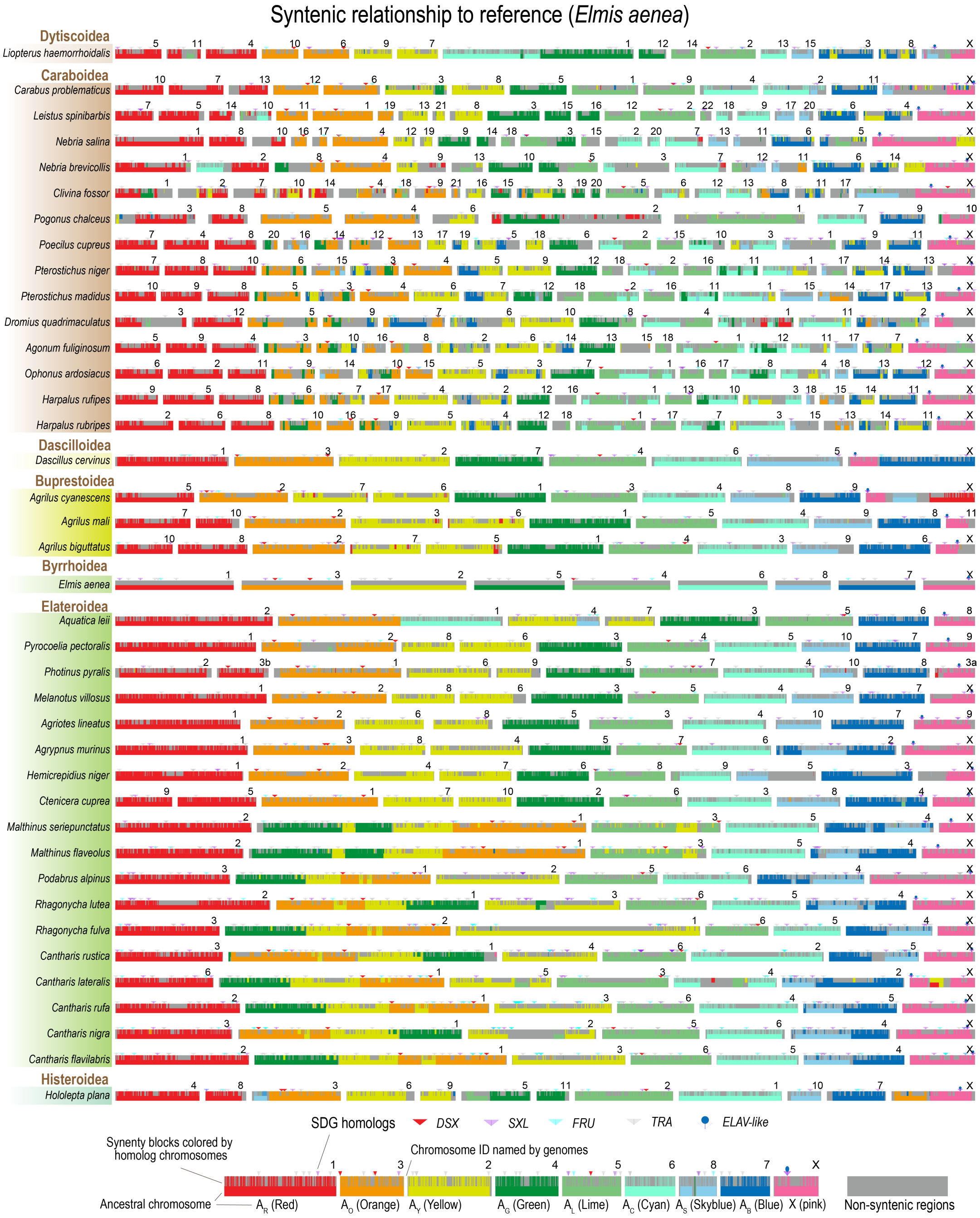

### (S1B) Superfamilies Staphylinoidea – Cleroidea:

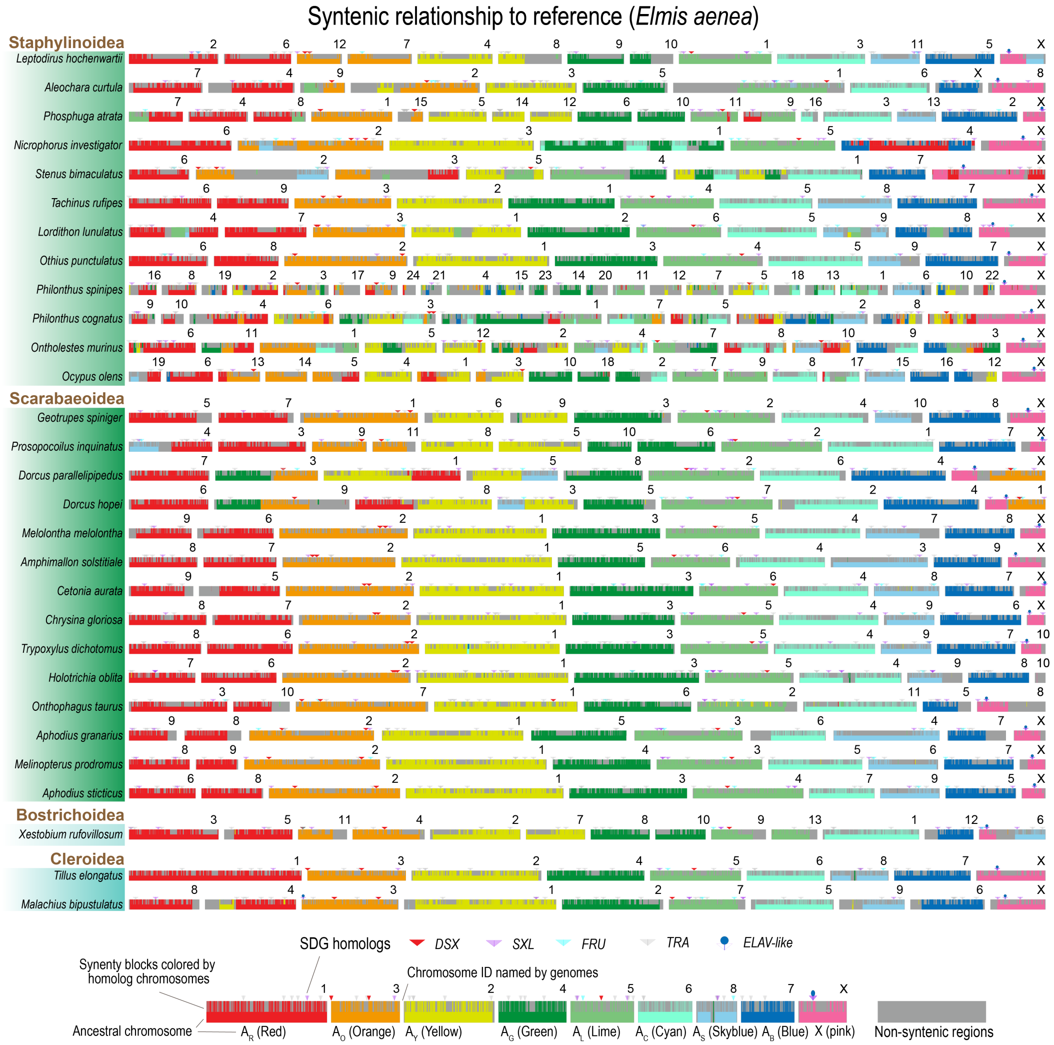

### (S1C) Superfamilies Coccinelloidea – Cucujoidea:

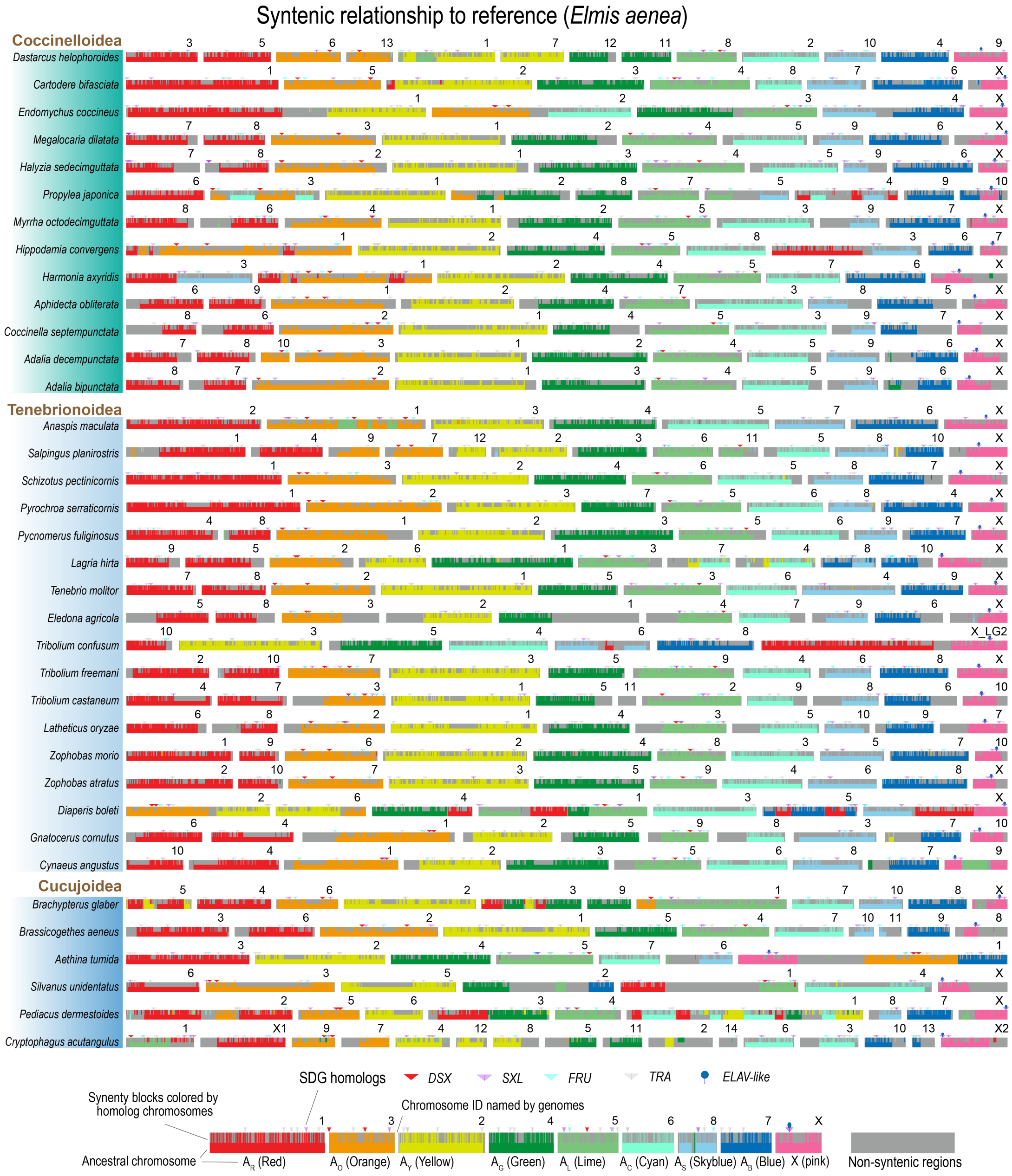

### (S1D) Superfamily Chrysomeloidea:

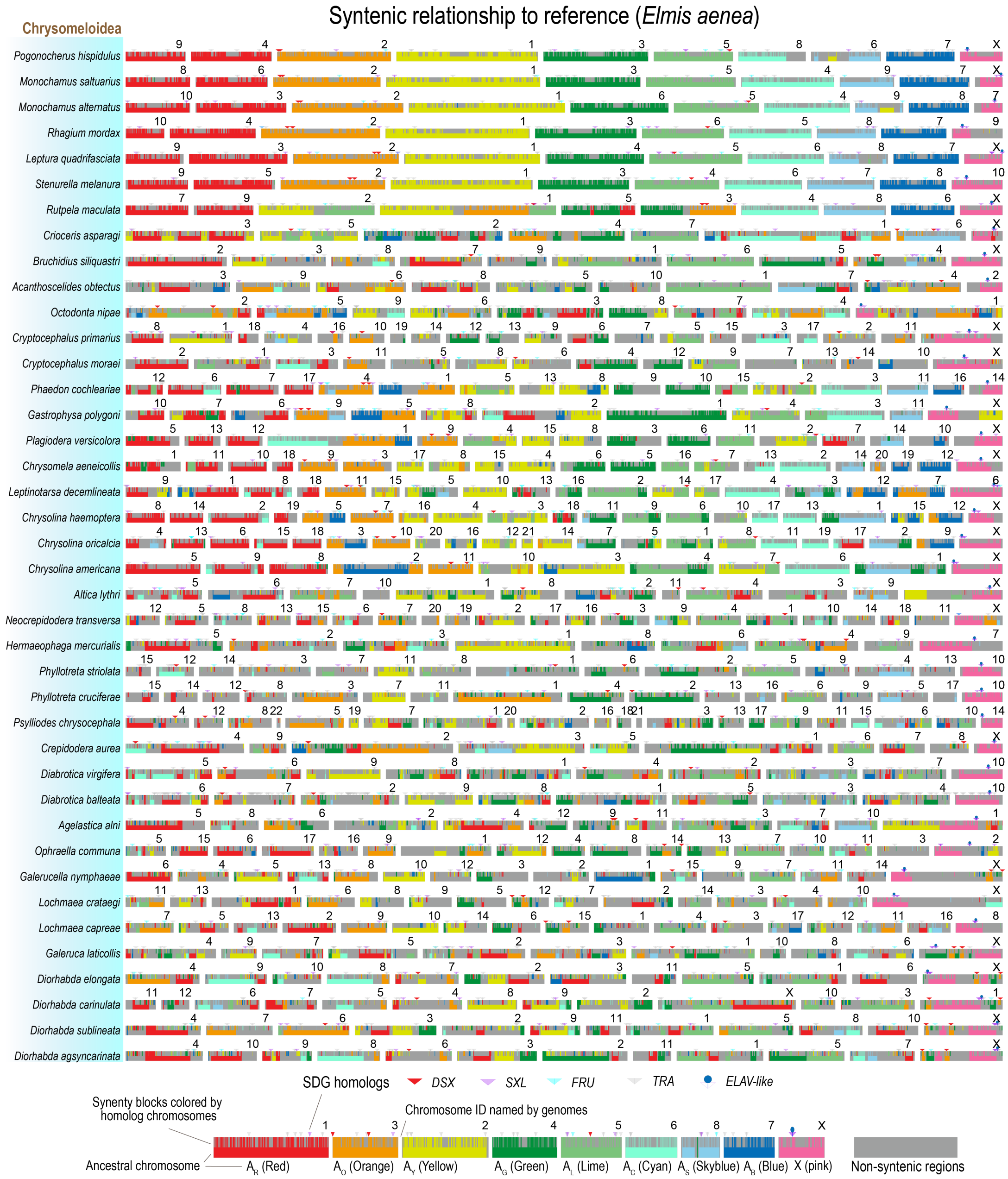

#### (S1E) Superfamily Curculionoidea:

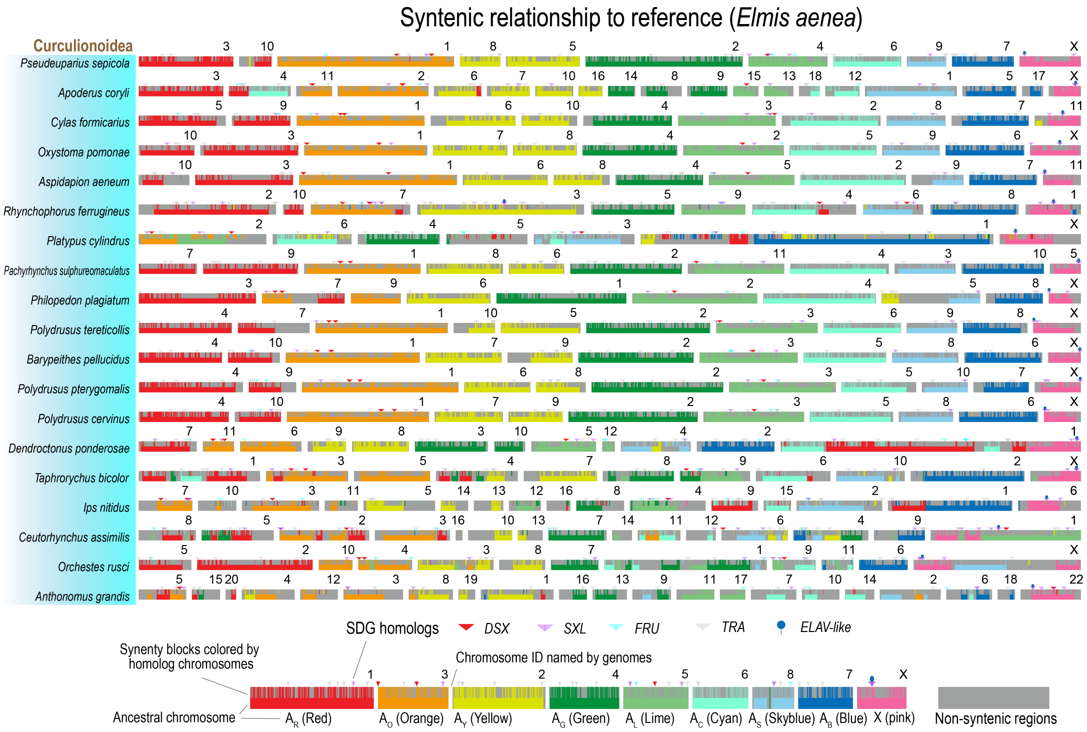

**Fig. S2.**

**Diverse chromosomal synteny patterns across beetles.** Pairwise synteny plots illustrating distinct patterns of chromosome evolution observed across beetle species. (**A**) Within *Agrilus* (*A. mali* and *A. cyanescens*; Buprestoidea): Autosomes are highly conserved. A neo-X chromosome in *A. cyanescens* originates from a fusion between *A. mali* chromosomes 10 and 11. (**B**) Within *Diabrotica* (*D. balteata* and *D. virgifera*; Chrysomeloidea): Chromosomal structure is completely conserved, with no inter-chromosomal rearrangements affecting either autosomes or the X chromosome. (**C**) Between *Dastarcus* and *Harmonia* (*D. helophoroides* and *H. axyridis*; Coccinelloidea): Most autosomes exhibit fusion or fission events, whereas the X chromosome remains structurally conserved. (**D**) Between *Rutpela* (*R.* maculata; Chrysomeloidea) and *Dascillus* (*D. cervinus*; Dascilloidea): Fusion and fission events are observed across both autosomes and the X chromosome. In *D. cervinus*, a neo-X chromosome is formed by fusion between an autosome and the ancestral X. (A–D) Coloured boxes indicate chromosome fissions/fusions.

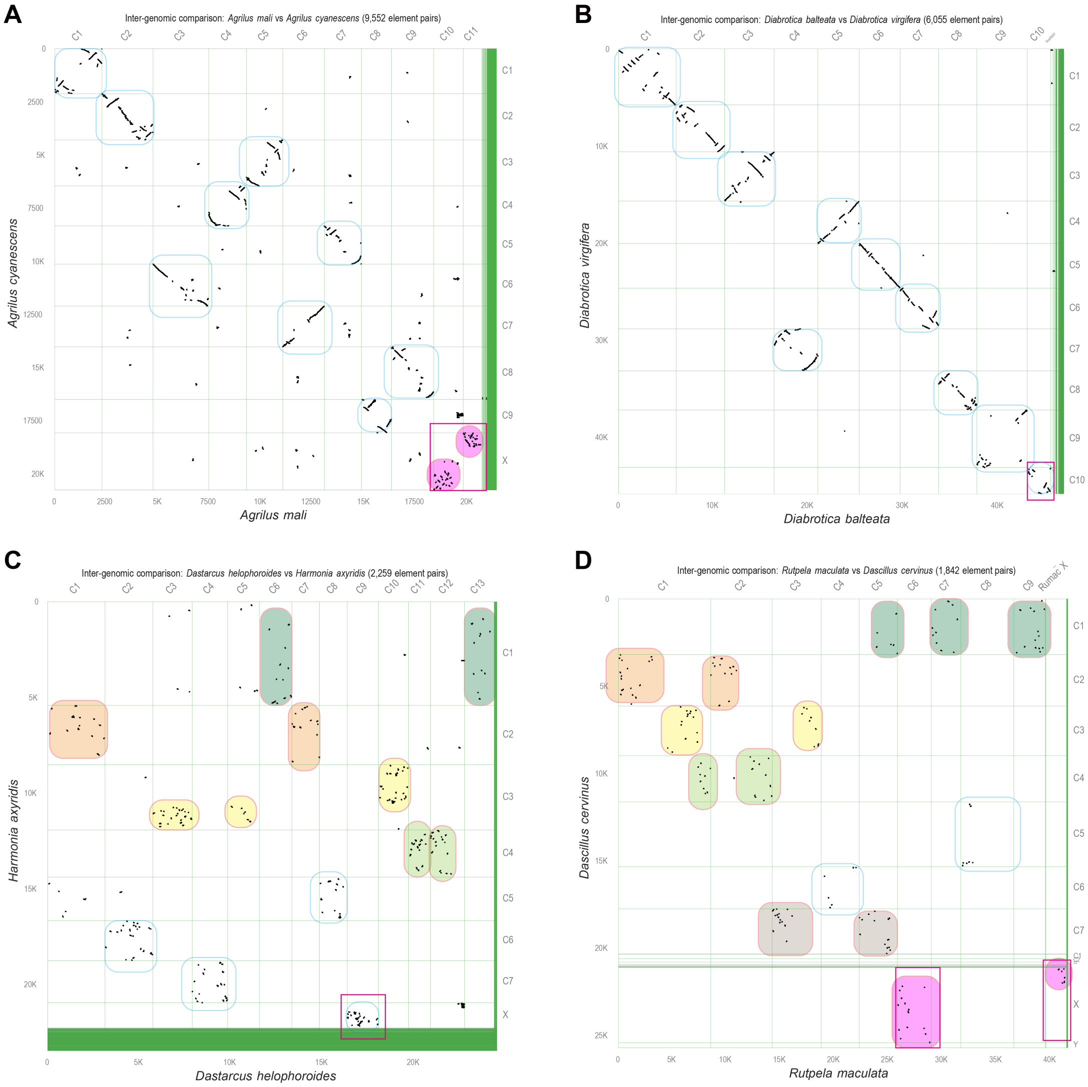

**Fig. S3.**

**Synteny across the Chrysomeloidea superfamily.** Phylogenetic tree inferred from 257 single-copy genes, with species arranged according to their evolutionary relationship. Horizontal brown bars represent the chromosomes of each species, with lengths drawn proportional to genome size (original chromosome labels shown in white). Syntenic blocks between species pairs are visualised as links: purple links indicate synteny on the conserved ancestral X chromosome region, whereas grey links indicate synteny on other genomic regions.

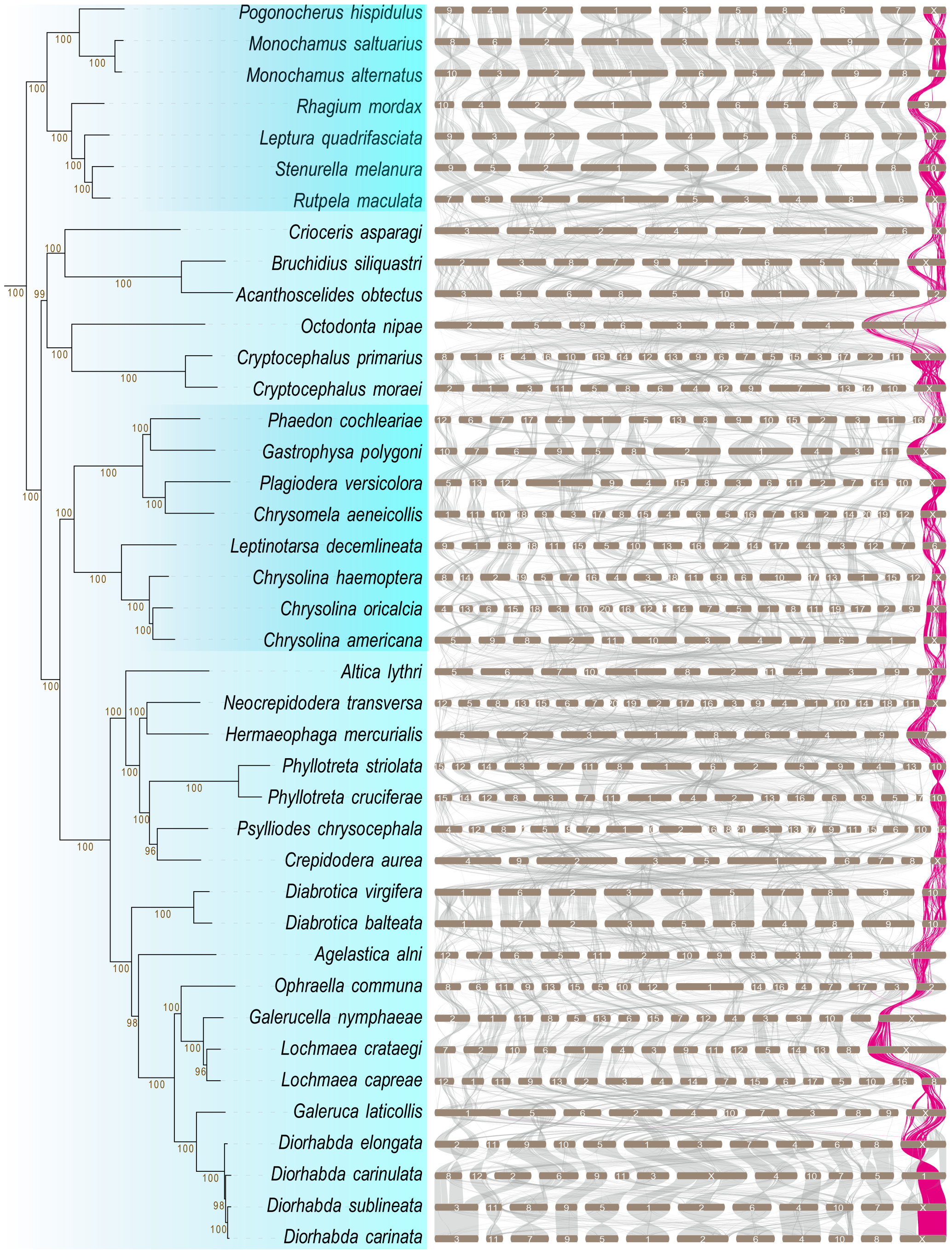

**Fig. S4.**

**Phylogenetic relationship and X chromosome evolution across 163 beetle species.** Phylogenetic species tree based on single-copy genes (branch support = 1 unless indicated). Also, shown are superfamily (gold) and species (black). Syntenic chromosome regions of each species relative to *Elmis aenea* reference chromosomes. The bar represents the X chromosome with the upper part showing syntenic blocks and the lower part the inferred reference chromosome regions (coloured by homology to the reference chromosomes). Positions of homologs of five key sex-determining genes (SDG homologs) are indicated above the bars by triangles. Also, shown is the proportion of the X chromosome composed of conserved (pink), translocated (black) and non-syntenic regions (grey).

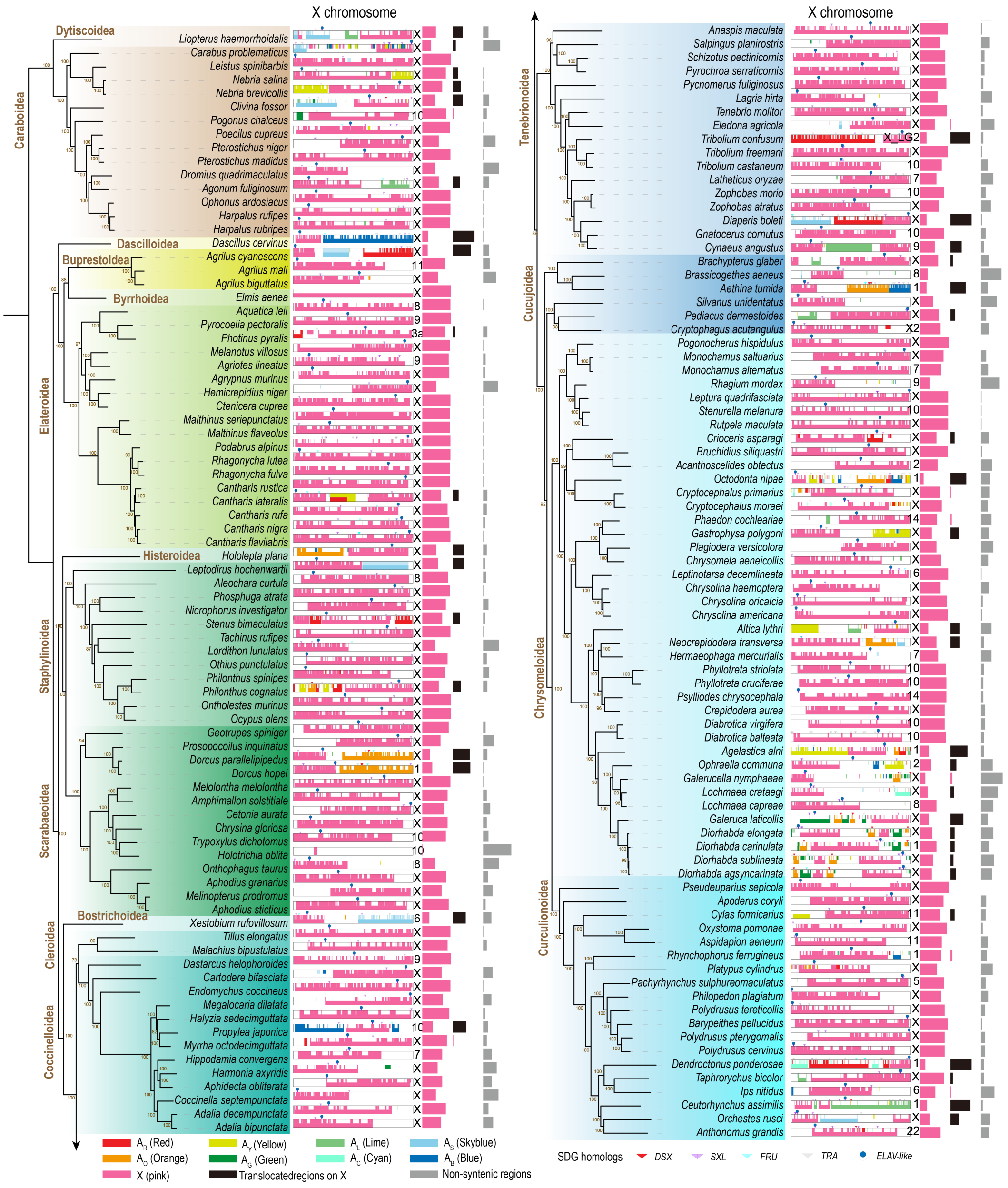

**Fig. S5.**

**Synteny network cluster profiling of X and Y chromosomes.** (**A**) Synteny network cluster relationships among X-linked homologs in 74 species, including species with neo-X chromosomes (51 species) and/or with assembled Y chromosomes (39 species). The X conserved region and some shared neo-X regions are indicated. (**B**) Synteny network cluster relationships among Y-linked homologs across 39 assembled Y chromosomes. (**C**) Chromosome length (black) and gene count (green) for the assembled Y chromosomes across the 39 species.

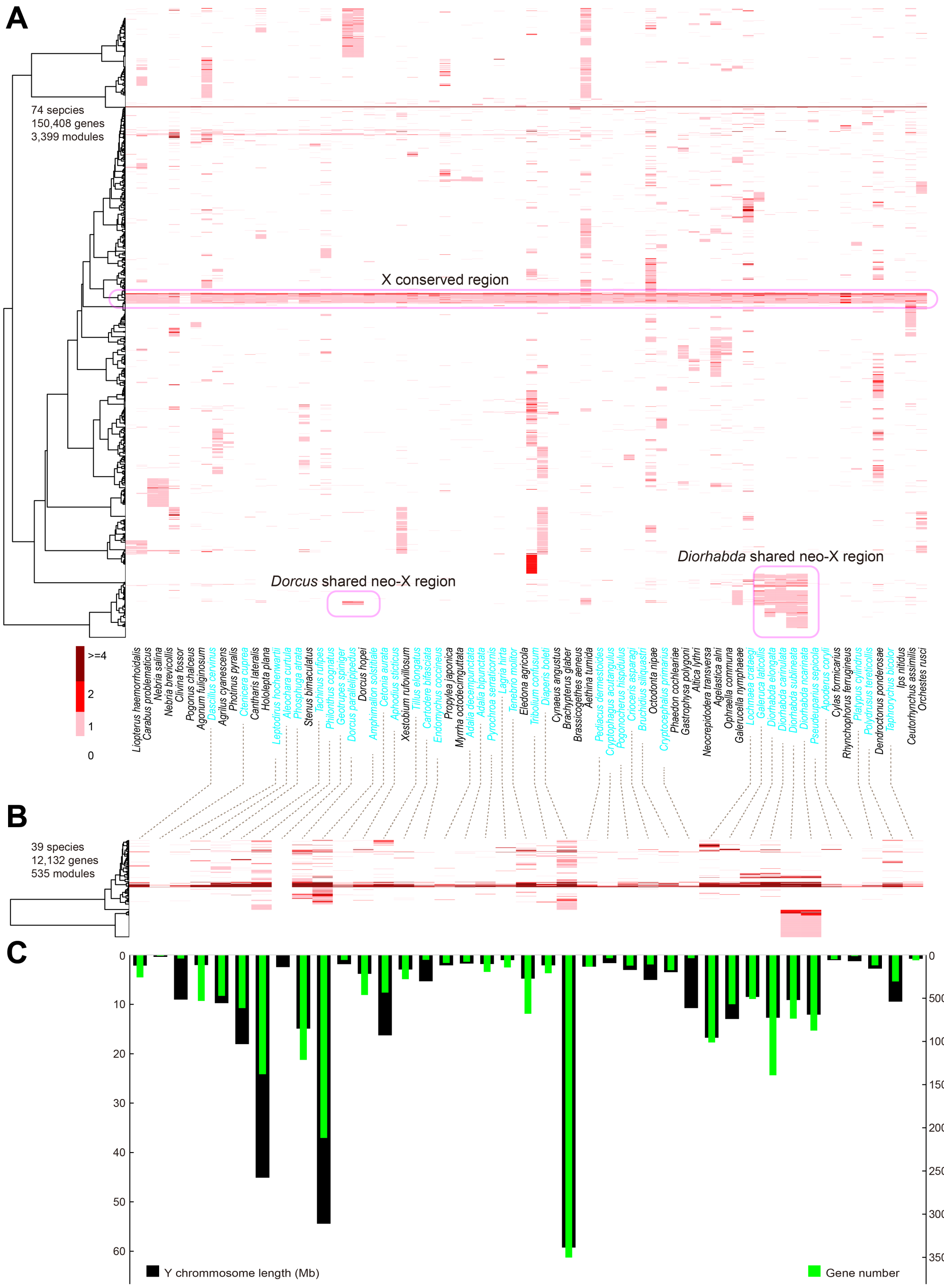

**Fig. S6.**

**Misidentified X chromosome in *Aleochara curtula* and misassembled X and Y chromosome in *Phosphuga atrata*.** *Nicrophorus investigator* is included as a reference, although this assembly lacks a Y chromosome. Graphs show pairwise syntenic blocks. (**A**) Absence of synteny between the X and Y chromosomes of *A. curtula* and *P. atrata*, indicating assembly errors. (**B**) Extensive synteny between the Y chromosome of *P. atrata* and the X chromosome of *N. investigator*, supporting misassembly of the Y chromosome in *P. atrata*. (**C**) Extensive synteny between chromosome 8 in *A. curtula* and chromosome X *N. investigator*, indicating that the X chromosome is misidentified in *A. curtula* (two other species also exhibited misidentified X chromosomes; Table S1). The of lack of syntenic blocks linking the assembled Y chromosome of *A. curtula* and the *N. investigator* assembly is consistent with a correctly assembled Y chromosome in *A. curtula*.

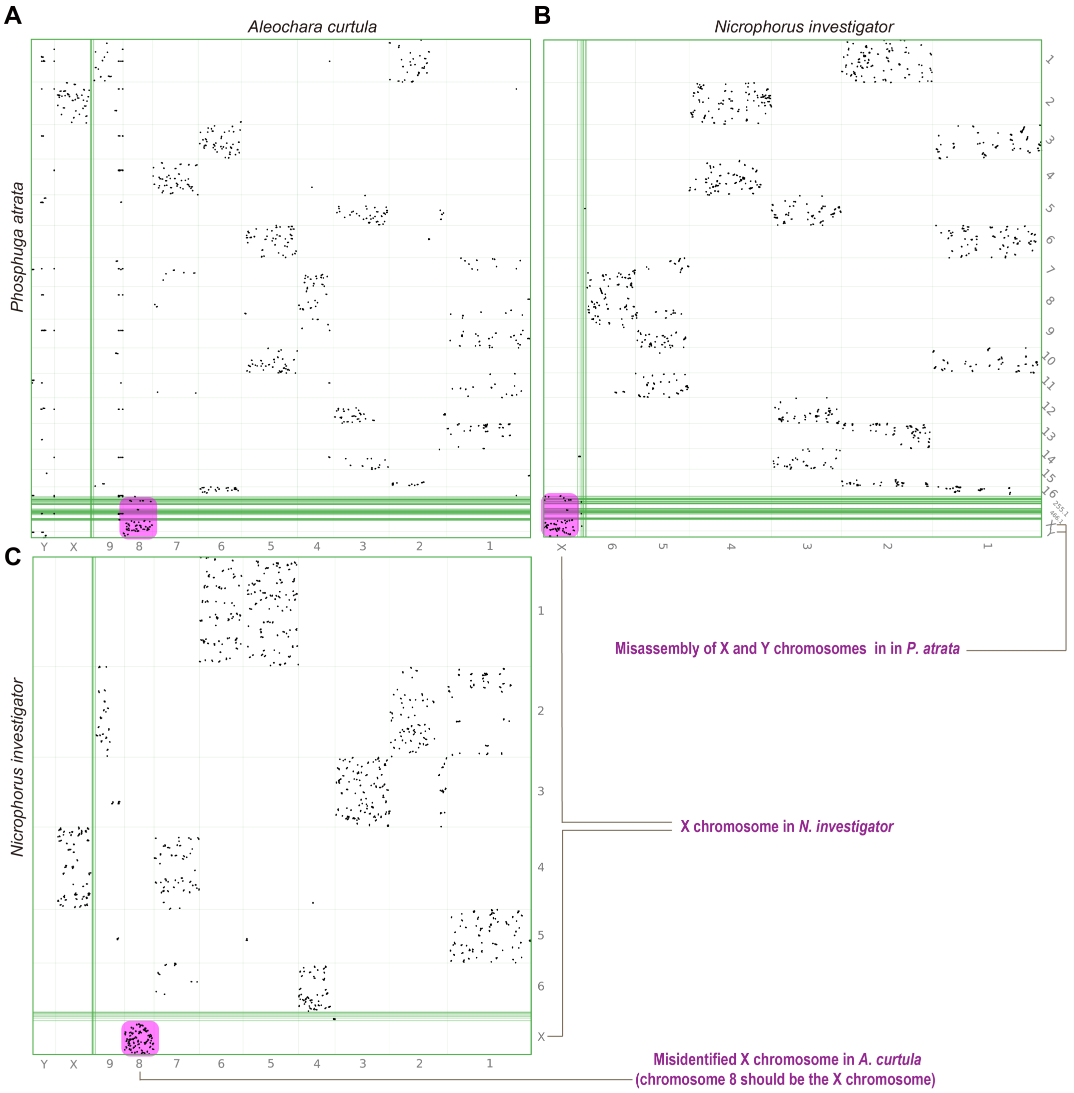

**Fig. S7.**

**Gene trees of homologs of three sex-determining genes**. (**A–C**) Gene trees of homologs of (**A**) *DSX*, (**B**) *TRA* and (**C**) *FRU* across 163 beetle genomes. Homologs located on the X chromosome are highlighted in pink. All gene models were predicted with the *AgSyn* pipeline (no filter applied).

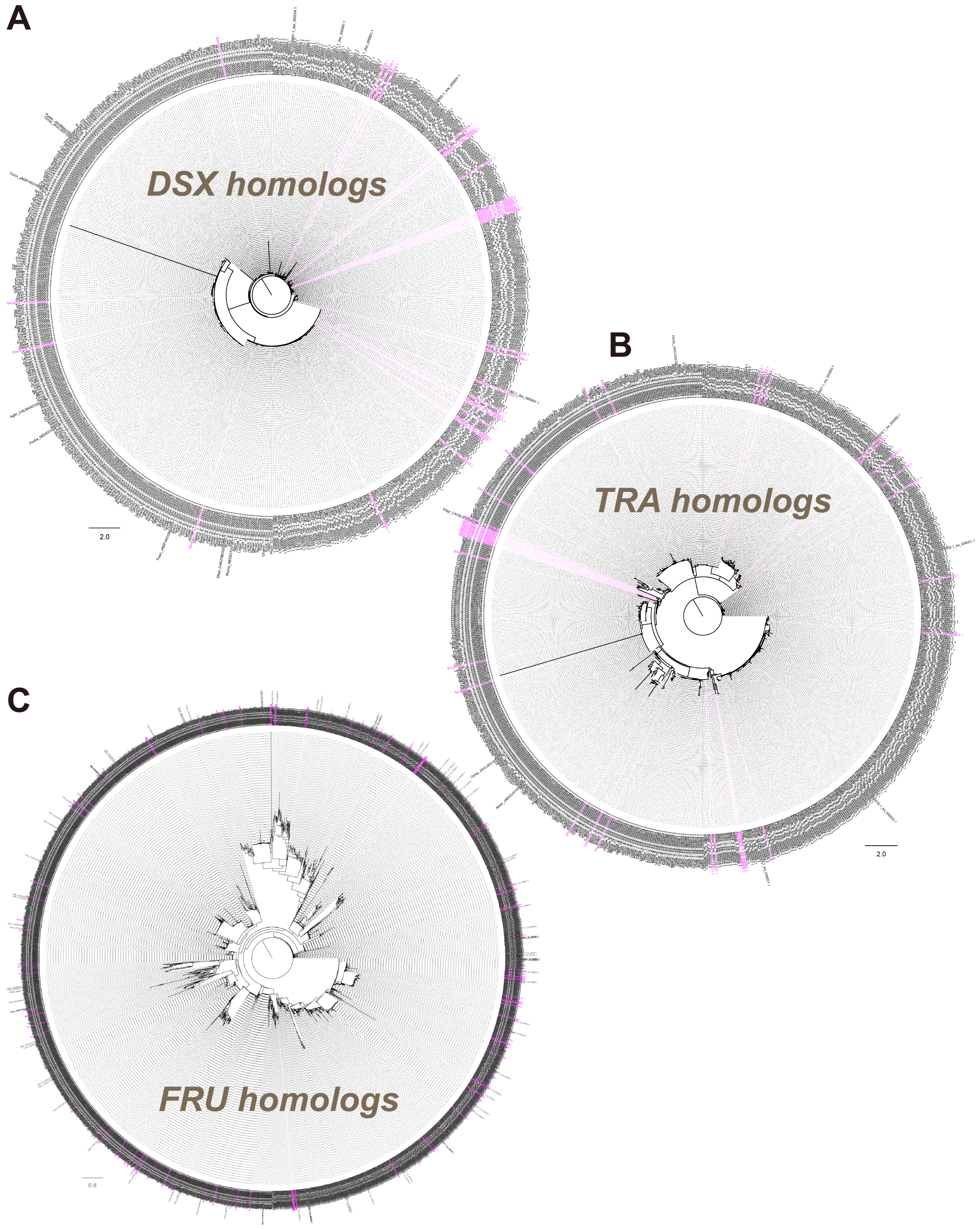

**Fig. S8.**

**Gene trees of *PABP-like*, *ELAV-like* and *SXL* homologs**. (**A–C**) Gene trees of homologs of (**A**) *PABP-like*, (**B**) *ELAV-like* and (**C**) *SXL* across up to 163 species. Genes located on the X chromosome are highlighted in pink. All genes were predicted using *AgSyn* and subsequently filtered to remove low quality sequences based on similarity to their reference genes (Fig. 5B), using *BLASTP* with the criteria: identity ≥ 80% and e-value ≤ 1e-120. (B) Green links indicate genes originating from the same species. (**D**) Chromosome location of *ELAV-like* in *Elmis aenea* where it is X-linked and *Pogonus chalceus* where it is located on an unanchored scaffold (Fig. 4B).

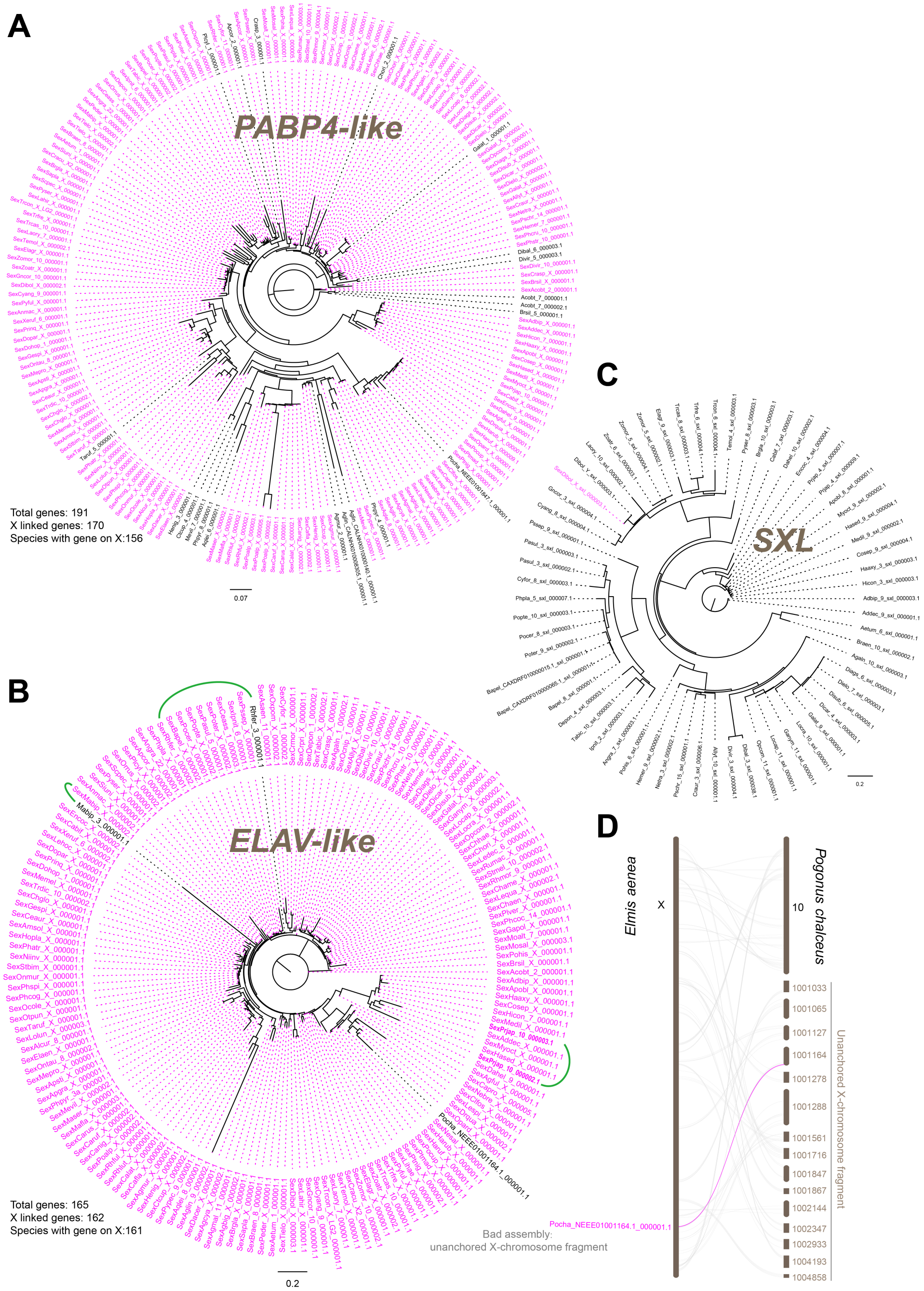

**Fig. S9.**

**Putative fission of the ancestral X in the $X_1X_2Y$ system of *Blaps rhynchoptera*.** (**A–B**) Pairwise synteny plots between chromosome-level genomes of *B. rhynchoptera* (NCBI accession number ASM4225770v1) and (A) a related species, *Tenebrio molitor*, and (B) the reference species *Elmis aenea*, illustrating the fission of ancestral X in *B. rhynchoptera* (purple box). (**C**) Pairwise synteny plots between *T. molitor* and *E. aenea* showing the conserved ancestral X region in these two species. (**D**) Links connecting syntenic blocks between *E. aenea*, *Tribolium castaneum*, *T. molitor* and *B. rhynchoptera*, further illustrating the X-chromosome fission in *B. rhynchoptera*.

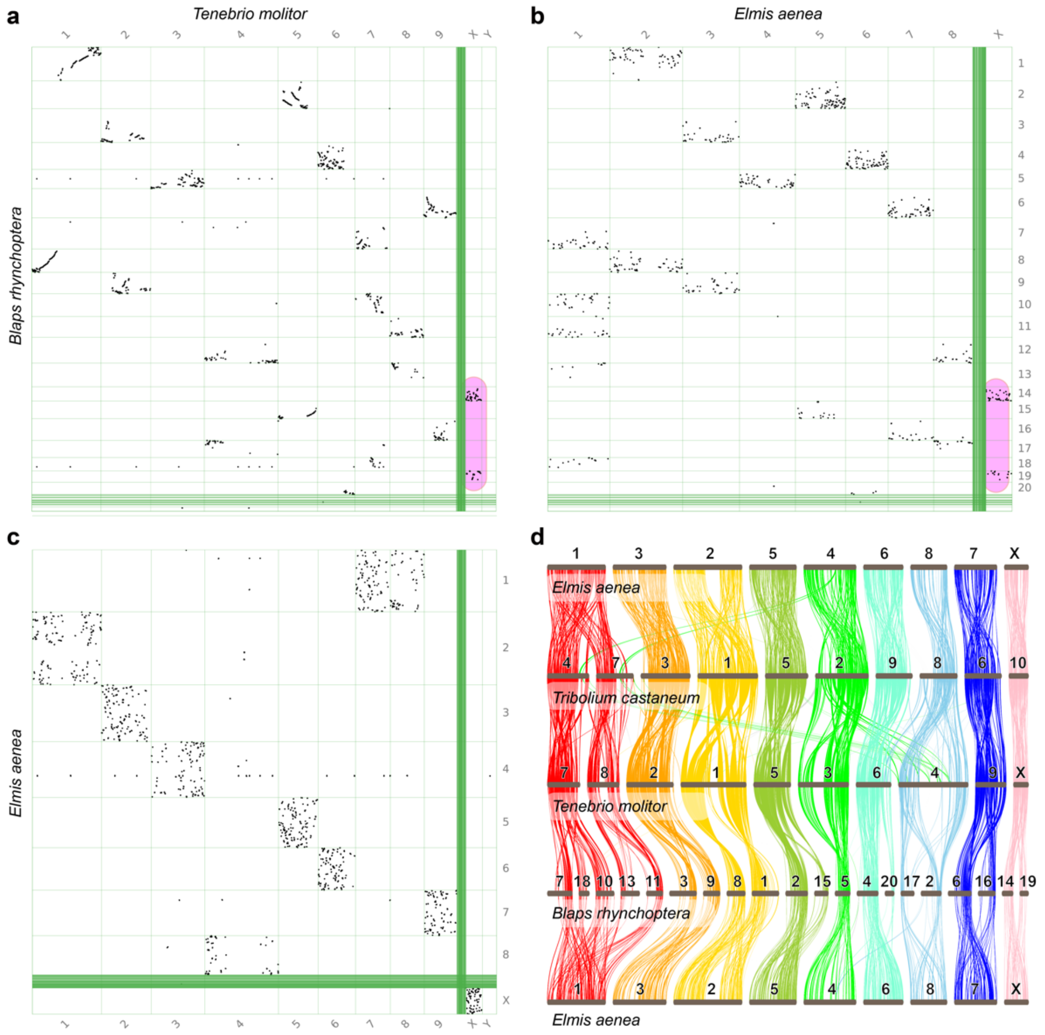

**Fig. S10.**

**Flowchart of the synteny analysis pipeline**. The pipeline integrates widely used tools (see Materials and Methods). Green boxes indicate input files (white text) and output files (black text).

Steps:

1. **Input genome assemblies.** Prepare and input chromosome-level or contig-level genome assemblies. When contig-level assemblies are used, anchor them to available chromosome-level genome (function *agysn.anchor*). Standardise chromosomes labels (1, 2, 3, …, X, Y). *We used chromosome-level assemblies of 163 beetle species (Table S1); thus, agysn.anchor was not used.*
2. **Split chromosomes.** To increase computational efficiency, identify simple repeats using *TANTAN* and split chromosomes at repeat regions into ~2 Mb fragments while retaining genome coordinates.
3. **Extract open reading frames (ORFs).** Predict and extract ORFs from split sequences using *ORF-FINDER*. Output file: ORFs.fa.
4. **Input reference protein sequences.** Provide high-quality reference protein sets from one or more well-annotated species and remove redundancies using *CD-HIT*. *We used protein sequences of five well-annotated beetle genomes and Drosophila melanogaster (see Materials and Methods).*
5. **Annotate assemblies.** Align ORFs to reference proteins using *DIAMOND* (*BLASTP*), remove ORFs lacking homologous evidence, and merge ORFs belonging to the same gene. ORFs within genes are marked as CDS. Output files: species.gff, gene models; species.cds, CDS sequences; species.pep, protein sequences. CDS and peptide files are generated from GFF using *gffread*.
6. **Obtain orthogroups (OGs).** Remove stop codons from protein sequences and assign OGs using *OrthoFinder*.
7. **Build phylogenetic tree.** Create multi-species alignments for single-copy OGs using *MAFFT* or *MUSCLE*, trim using *TrimAl* and infer gene trees using *IQ-TREE*. Gene trees are then summarised to a species tree using *ASTRAL*. *We used 1878 single-copy genes from 16 species and 515 single-copy genes from 163 species to construct beetle phylogenies (Figs. 1A–2A).*
8. **Analyse synteny.** Identify synteny blocks and links to a reference genome using *JCVI*, with annotated files (*gff*, *cds* and *pep*) as input. Ordering chromosomes (function *agsyn.sort*). Visualise global syntenic relationships (function *agsyn.syn*; Fig. S1A–E). Synteny between two bird genomes (*Myiopsitta monachus* and *Gallus gallus*; NCBI accession numbers GCA_017639245.1 and GCA_024206055.2) is shown as an example (coloured insert; syntenic links coloured according *Gallus gallus* chromosomes; dotplot and reconstruction of chromosome configuration generated with *WGDI*). *Beetle synteny outputs are shown in Figs. 1–3 and Figs. S1–S4.*
9. **Identify sex chromosome-specific OGs.** Identify and extract OGs linked to sex chromosomes (X and Y), scaffolds and autosomes (function: *agsyn.sex*). Then annotate function of all OGs (function: *agsyn.agsx.func*): align OGs to well-annotated reference genomes via *BLASTP* to determine the function of specific genes. Finally, display the selected sex chromosome-linked genes (Fig. 4A–B). *We used the reference genomes of Aethina tumida and Diorhabda carinulata.*
10. **Identify sex-determining gene (SDG) homologs.** Identify homologs of key SDGs from well-annotated genomes (function: *agsyn.genewise2*), which employs *TBLASTN*, *SOLAR*

and *GeneWise2*. *We used four key SDGs from eight well-annotated genomes (see Materials and Methods).*

11. **Build gene trees.** Align SDG candidate sequences with *MAFFT* or *MUSCLE*, trim with *TrimAl*, and infer SDG gene trees with *IQ-TREE*. *We built gene trees of homologs of DSX, FRU, SLX and TRA (Fig. 5A; Figs. S7–S8).*
12. **Confirm ancestral sex chromosome.** Having identified candidate sex chromosomes from synteny (step 9), confirm them by examining whether SDG homologs (step 11) occur almost exclusively on sex chromosomes and rarely on autosomes across species.
13. **Additional downstream analyses.** The annotated output generated above can be used for further analyses of synteny, phylogeny or gene families using tools such as *WGDI*, *Infomap* and other pipelines for chromosome-evolution studies. Dotplot and reconstruction of chromosome configuration of *Myiopsitta monachus* and *Gallus gallus* generated with *WGDI*, and synteny network cluster relationship among X-linked genes in 31 beetle species generated with *Infomap*, are shown as examples (coloured inserts). *The present study includes an analysis of the synteny network cluster relationship across X and Y chromosomes (Fig. S5A–B).*

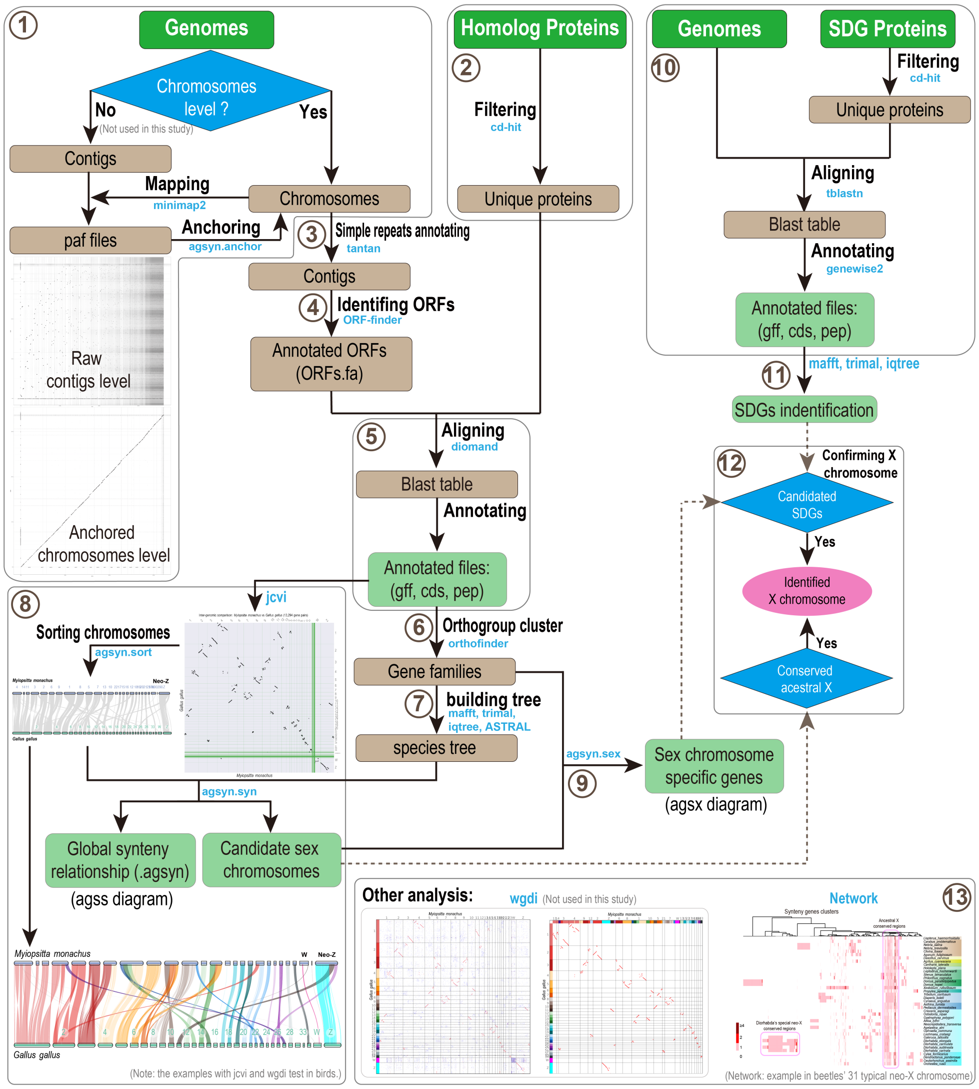

**Fig. S11.**

**Running time of the pipeline.** Runtime estimates based on 12 beetle genomes ranging 372 Mb to 2533 Mb. On average, the pipeline annotates ~120Mb genome sequence per 10 minutes, corresponding to a throughput of ~0.5 Gb per CPU-hour.

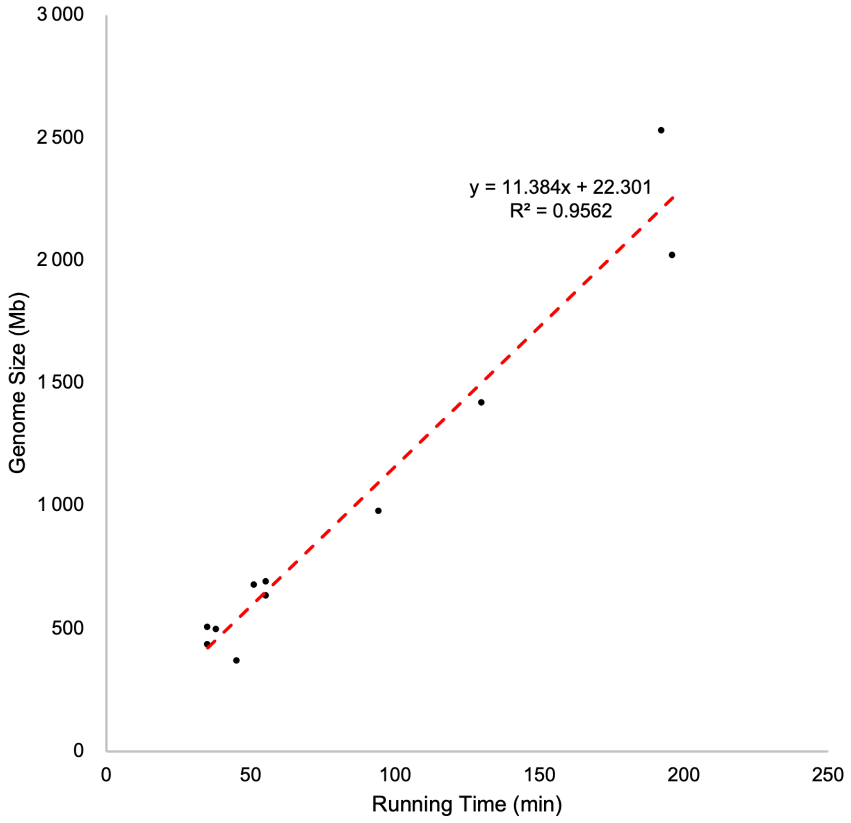
